# Actomyosin contractility in olfactory placode neurons opens the skin epithelium to form the nostril

**DOI:** 10.1101/2022.07.15.500210

**Authors:** Marion Baraban, Clara Gordillo Pi, Isabelle Bonnet, Jean-François Gilles, Camille Lejeune, Mélody Cabrera, Florian Tep, Marie Anne Breau

## Abstract

Despite their barrier function, epithelial layers can locally lose their integrity to create physiological openings during morphogenesis. The cellular and molecular mechanisms driving the formation of these epithelial breaks are only starting to be investigated. Here, we studied the formation of the zebrafish nostril (the olfactory orifice), which opens in the skin epithelium to expose the olfactory neurons to external odorant cues. Combining live imaging, drug treatments, laser ablation and tissue-specific functional perturbations, we demonstrate that the formation of the orifice is driven by a mechanical interplay between the olfactory placode neurons and the skin: the neurons pull on the overlying skin cells in an actomyosin-dependent manner, thus triggering the opening of the orifice. This work unravels an original mechanism to break an epithelial sheet, in which an adjacent group of cells instructs and mechanically assists the epithelium to induce its local rupture.

## Introduction

Epithelial barriers can disrupt their architecture to allow the formation of transient or stable gaps required for organ morphogenesis. The peripodial layer surrounding the *Drosophila* leg imaginal disc is for instance known to break before being removed through a curling process to facilitate leg eversion (Proag et al., 2019; Fouchard et al., 2020). A transient and local opening occurs at tricellular junctions of the *Drosophila* follicule epithelium and is essential for the import of yolk proteins in the oocyte (Isasti-Sanchez et al., 2021; Row and Deng, 2021). The mechanisms underlying the opening of epithelial holes have been attracting a growing interest in the field of epithelial morphogenesis (Bonfanti et al., 2022). These gaps open in the plane of the epithelium, which differs from the well-studied formation of enclosed lumens on the apical side of epithelial cells (Sigurbjörnsdóttir et al., 2014; Torres-Sanchez et al., 2021), and of tubular organs through invagination of the epithelium (Gilder and Röper, 2014).

Here, we focused on the development of the olfactory organ in zebrafish. The emergence of sensory functions often requires the formation of breaks through epithelial barriers covering the external or internal embryonic surfaces, allowing the exposure of sensory cells to chemical and mechanical cues from the oustide. This is illustrated by the lateral line mechanosensory neuromasts and the taste buds, in which a pore forms above the sensory cells, through yet unknown mechanisms, within the skin and the tongue epithelia, respectively (Hansen et al., 2002; Gleason et al., 2009; Barlow, 2015; Dow et al., 2018; Hino et al., 2022). Similarly, in the zebrafish olfactory organ, the sensory neurons are directly exposed to the outside through a gap in the skin epithelium, the olfactory orifice (Hansen and Zeiske, 1993; Cheung et al., 2021). This hole in the skin is essential for the setting of the olfactory function as it prefigures the future nostril allowing the detection of odorant cues by the sensory neurons (Hansen and Zeiske, 1998; Reiten et al., 2017; Cheung et al., 2021), yet the cell behaviours and molecular mechanisms involved in its formation are totally unknown. Previous studies reported the presence of a rosette of neurons located in the dorsal (apical) part of the olfactory placode, organising around the olfactory orifice (Whitlock and Westerfield, 1998, Yoshida et al., 2002, Miyasaka et al., 2005; Madelaine et al., 2011; Kress et al., 2015). These neurons show a circular organisation and extend dendrites towards the orifice. Based on their morphology, their superficial localisation and their expression of olfactory receptor genes, they were identified as precursors of the sensory neurons detecting the odor cues from the environment, as opposed to more ventral (or basal), dendrite-less neurons proposed to represent pioneer neurons (Whitlock and Westerfield 1998; Sato et al., 2005; Saxena et al., 2013). However, the link between the placodal rosette and the formation of the olfactory orifice has never been characterised.

We took advantage of this system to investigate the biomechanical factors driving epithelial hole opening *in vivo*. Using live imaging, we observed that the olfactory orifice opens through the local loss of cell/cell contacts in the skin, above the centre of the placodal rosette. We combined tissue-specific functional perturbations, drug treatments and laser ablation to show that mechanical interactions between olfactory placode neurons and the overlying skin epithelium play a crucial role in the formation of the olfactory orifice. Our results support a scenario in which the apical tips of the olfactory placode neurons forming the rosette pull on overlying skin cells, thereby inducing the local loss of homotypic cell/cell contacts and the hole opening. This work highlights a previously unknown mechanism controlling the physiological opening of an epithelial sheet *in vivo*.

## Results

### The olfactory placode is covered by a monolayer of peridermal skin cells prior to the opening of the olfactory orifice

From images in the literature we identified that the olfactory orifice opens between 24 and 48 hpf (hours post-fertilisation) (Araya et al., 2014; Piotrowski and Nüsslein-Volhard, 2000). To determine the precise developmental stage at which the olfactory orifice opens, we performed a time-course analysis using *Tg(-8.0cldnb:LY-EGFP)* transgenic embryos in which cells of the olfactory placode and the periderm, *i.e.* the most superficial skin layer, are labelled with membrane-targeted GFP (Haas and Gilmour, 2006) (Figure 1A). Up to 26 hpf, no orifice had formed and the skin epithelium constituted an intact physical barrier covering the olfactory neurons. From 28 hpf, the skin layer started to break above the olfactory placode in some embryos, leading to the formation of an orifice in 79% of the embryos at 32 hpf, and in 100% of the embryos at 34 hpf (Figure 1A’). Thus, the process of opening is not synchronous in same-stage embryos and occurs gradually between 28 and 34 hpf (Figure 1A, A’) (n = 123 embryos, 3 independent experiments).

**Figure 1.**
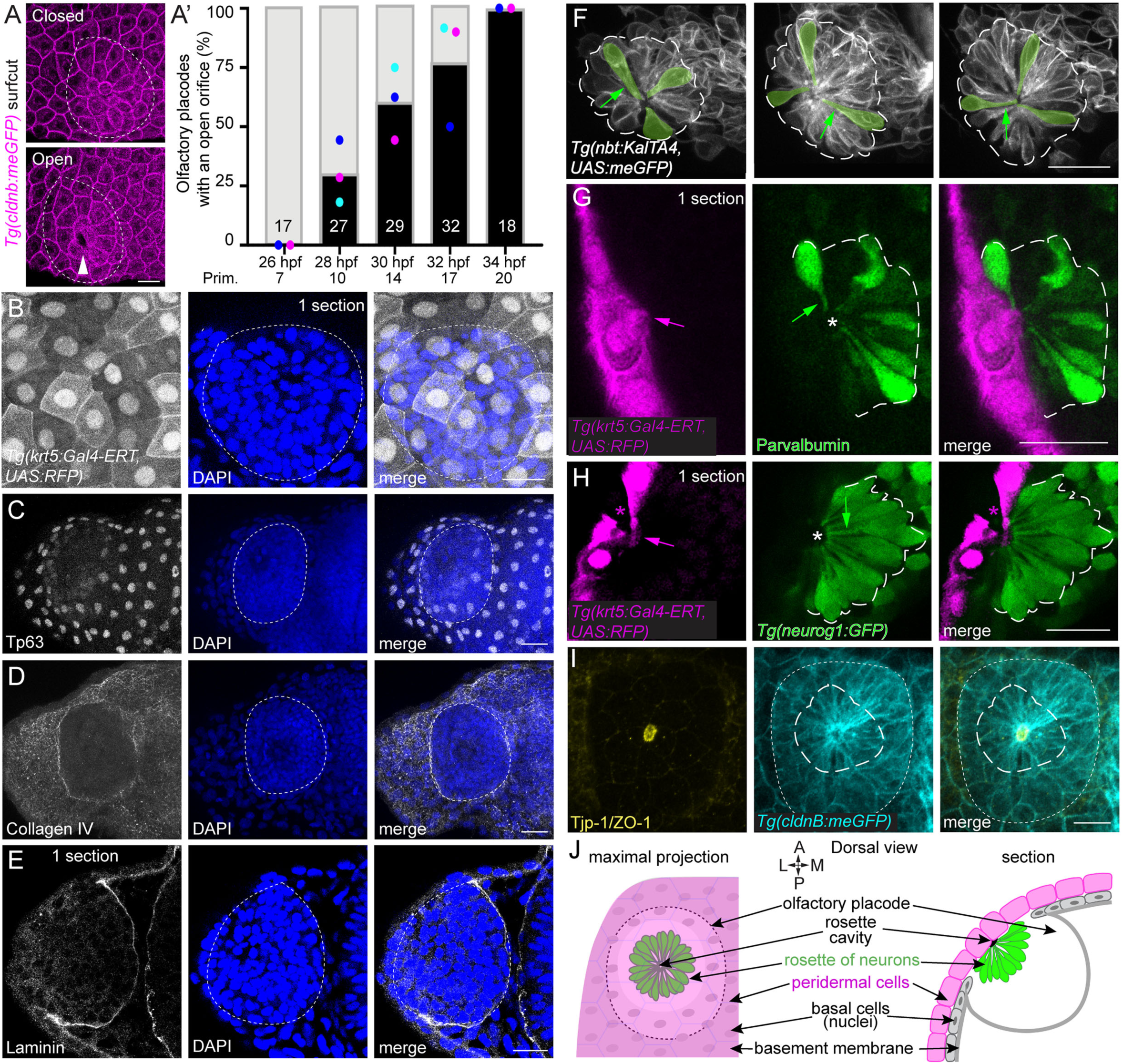
Interaction between the skin epithelium and the olfactory neurons before the opening of the olfactory orifice. All images shown in this Figure are dorsal views, oriented anterior to the top, and lateral on the left. **A**. Images of 32 hpf *Tg(cldnb:meGFP)* embryos in which the olfactory orifice is closed (top) versus open (bottom). The image J plugin SurfCut (Erguvan et al., 2019) was used to extract the periderm layer above the olfactory placode and determine if an orifice was present (see Methods). The contour of the olfactory placode, seen in deeper z-positions, is indicated by white dotted lines. **A’**. Graph showing the percentage of olfactory placodes with an open orifice at various stages between 26 and 34 hpf. To precisely stage the embryos, we used the position of the migrating posterior lateral line primordium (Kimmel et al., 1995), which is also labelled in *Tg(cldnb:meGFP)* embryos (Haas and Gilmour, 2006). The position (in somites) of the lateral line primordium is also indicated for each stage. The color dots indicate the results for 3 independent experiments (n = 123 embryos in total, the number of embryos analysed at each stage is indicated on the columns of the graph). **B.** Images of a *Tg(krt5:Gal4-ERT, UAS:RFP*) double transgenic embryo at 26 hpf, showing the periderm that expresses RFP (grey), and the DAPI nuclear labelling (blue) allowing to visualise the position and morphology of the olfactory placode (dotted line) below the skin. The placode is covered by a continuous monolayer of peridermal cells at this stage. Note that on fixed embryos carrying the *Tg(krt5:Gal4-ERT, UAS:RFP*) transgenes, we observed that, for unknown reasons, the RFP signal is stronger in the nucleus that in the cytoplasm of peridermal cells. **C.** Immunostaining for the nuclear basal cell marker Tp63 (grey) and DAPI staining (blue) at 26 hpf showing a basal cell nuclei-free area above the dorsalmost region of the olfactory placode. The dotted lines indicate the contour of the olfactory placode. **D, E.** Immunostainings for Collagen IV (D) and Laminin (E) (grey) and DAPI staining (blue) at 26 hpf showing the absence of these basement membrane components at the interface between the skin and the olfactory placode. The placode is indicated by dotted lines. **F.** Images of 27 hpf embryos carrying the *Tg(nbt:KalTA4, UAS:meGFP)* transgenes labelling a subgroup of the rosette neurons. The membrane labelling allows to visualise the radial organisation and the pear-shaped morphology of the rosette neurons, which extend their dendrites (green arrows) towards the centre of the rosette. **G, H.** Interaction between the monolayer of peridermal cells (labelled with *Tg(krt5:Gal4-ERT, UAS:RFP)* in magenta) and the olfactory neurons (labelled in green with Parvalbumin immunostaining in G, and *Tg(neurog1:GFP)* in H), assembled into a rosette just before orifice opening, at 29 hpf. The white asterisk indicates the hemispheric cavity forming in the centre of the rosette and referred to as the rosette cavity. The green arrows point to dendrites. The magenta asterisk indicates a peridermal cell which does not express the *Tg(krt5:Gal4-ERT, UAS:RFP)* transgenes (likely due to variegation) but is present above the rosette cavity (see Video S1). The magenta arrows show examples of deformations or cytoplasmic protrusions of skin cells directed towards the rosette cavity. **I.** Images of a 26 hpf *Tg(cldnb:meGFP)* embryo immunostained for GFP (blue) and for the tight junction protein ZO-1 (or Tjp-1) (yellow), showing the accumulation of ZO-1 at the apical tips of the dendrites of the rosette neurons, and also at the cortex of peridermal cells. ZO-1 staining at the border of the rosette cavity was thus used as a readout of the size and shape of this structure (see Video S2). **J.** Schematic view showing the system before the opening of the olfactory orifice in a dorsal view, in maximal projection and in one z-section, summarising the observations presented in Figure 1. Scale bars: 20 µm except C and D: 25 µm.

At these stages, the embryonic skin of zebrafish embryos is composed of the superficial peridermal cells covering the basal cells, which themselves secrete a deeper layer of basement membrane extracellular matrix (ECM) (O’Brien et al., 2012). To analyse whether the stratified architecture of the skin is conserved above the olfactory placode, we used a combination of transgenic lines and antibodies to reveal the skin layers at 26 hpf, a few hours before the opening of the orifice. Using *Tg(krt5:Gal4-ERT, UAS:RFP*) double transgenic fish, in which the periderm expresses RFP (Akerberg et al., 2014), and DAPI labelling to visualise the olfactory placode (which appears as a compact cell cluster), we confirmed that the placode is covered by a continuous peridermal layer at this stage (Figure 1B). Surprisingly, the basal cell nuclei did not entirely cover the olfactory placode, leaving an apparent basal cell-free area in its dorsalmost region (Figure 1C). In addition, Laminin and Collagen IV, two ECM components of the basement membrane, were present around the basal and lateral regions of the placode, but absent from the interface between dorsal placode cells and the periderm (Figure 1D, E). Thus, before the opening of the olfactory orifice, the skin overlying the olfactory placode is exclusively composed of a monolayer of peridermal cells, which will be referred to as skin cells. We next analysed the organisation of the olfactory placode neurons and their interaction with the skin cells.

### The rosette of olfactory placode neurons is tightly apposed to the skin cells before the opening onset

Previous work described the assembly of a rosette of neurons localised in the dorso-lateral region of the olfactory placode (Whitlock and Westerfield, 1998, Yoshida et al., 2002, Miyasaka et al., 2005; Madelaine et al., 2011; Kress et al., 2015). We analysed the cellular organisation in the olfactory placode and confirmed the presence of a rosette of neurons from 25 hpf onwards, visualised with a variety of neuron-specific transgenic lines and antibodies against neuronal markers (Figure 1F-H, for 3D visualisation the z-stack of the Figure 1H panel is shown in Video S1). Right before the opening, at 26-27 hpf, the dorsal rosette of placodal neurons was observed just adjacent to the monolayer of peridermal cells covering the olfactory placode (Figure 1G-H), with the apical tips of the dendrites being in close contact with the skin. In addition, we found that a hemispheric cavity is present in the centre of the rosette (Figure 1G, H and Video S1) and is lined by the ZO-1 (Zonula Occludens-1)-labelled apical surfaces of the dendrites from the sensory neurons (Figure 1I and Video S2). This structure prefigures the future hollow, cup-shaped cavity of the nose pit in larvae (Reiten et al., 2017) and will be referred to as the rosette cavity. Altogether, our data show that the peridermal epithelial layer is juxtaposed to the rosette of placodal neurons before the opening of the olfactory orifice (Figure 1J). We then set out to analyse the skin cell behaviours driving the epithelial opening.

### The olfactory orifice opens through the loss of contacts between skin cells

To record the dynamics of orifice formation and analyse the associated skin cell behaviours, we performed live imaging on transgenic embryos in which fluorescent proteins are specifically expressed in peridermal skin cells (n = 44 movies from 14 independent experiments). Between 28 and 32 hpf, in all the movies, we detected the formation of a small, discrete hole in the skin layer, which opened through the local loss of cell-cell contacts between peridermal cells (Figure 2A-C, Videos S3 and S4). We found extremely rare cases of cell death events associated with the opening of the orifice (only 2 out of 44 cases, Figure 2C). Cell divisions were seen during and at the location of orifice formation in 9 out of 44 movies (Figure 2C). This proportion (20%) does not exclude a potential contribution of cell division in some embryos, however it seems not to be systematically associated with orifice formation. These observations led us to exclude cell death or division as the main drivers of the opening of the olfactory orifice, which relies mainly on the local loss of adhesion between skin cells, occuring in 100% of the cases (Figure 2C). The opening speed was different between embryos from the same clutch, showing the variability in the dynamics of the process (Figure 2D).

**Figure 2.**
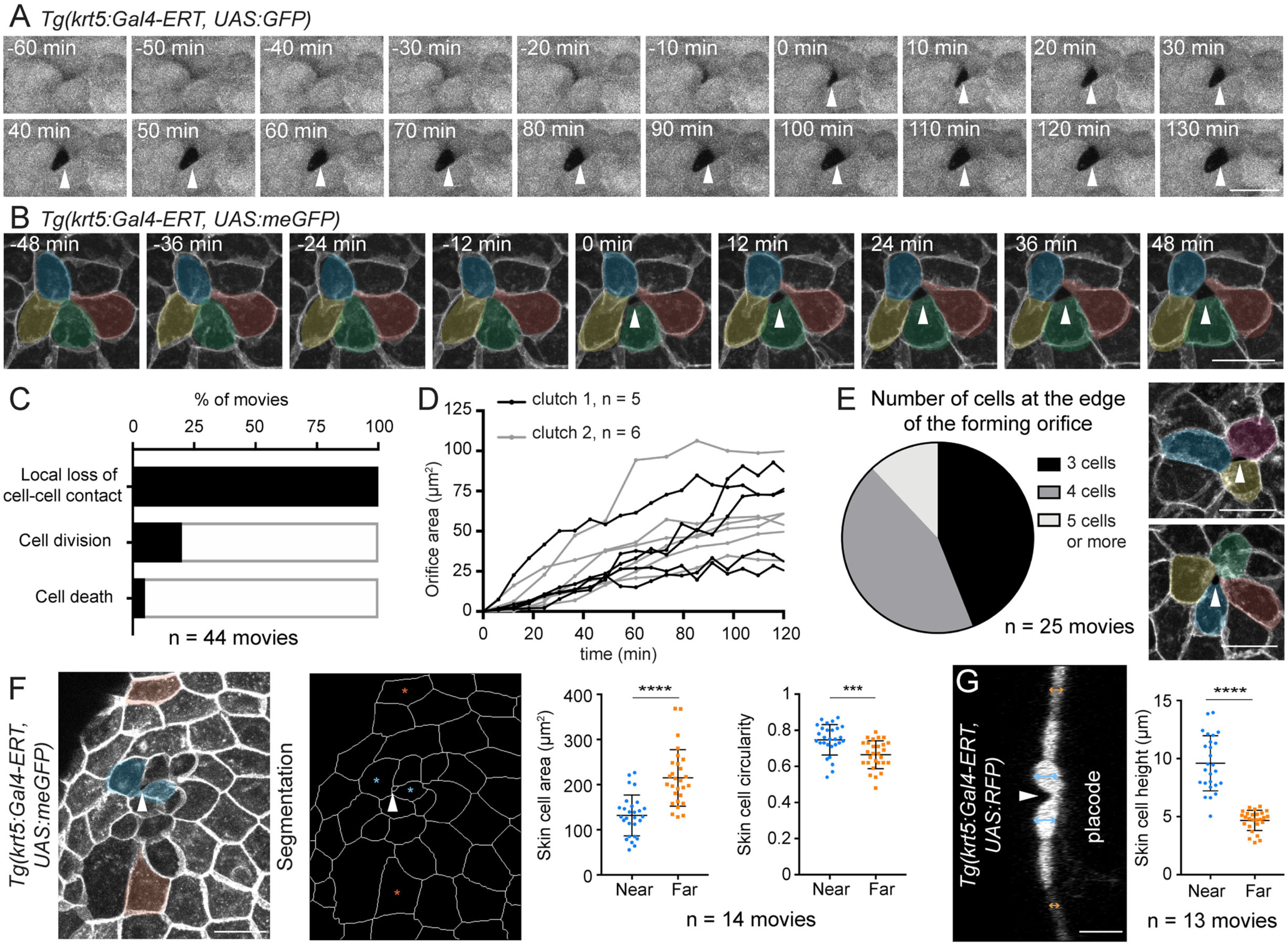
**Dynamic behaviour of the skin cells during the opening of the orifice**. All images shown in this Figure are dorsal views, oriented anterior to the top, and lateral on the left. **A.** Live imaging on a *Tg(krt5:Gal4-ERT, UAS:GFP)* embryo showing the opening of the olfactory orifice (indicated with a white arrowhead) through the progressive loss of contacts between peridermal skin cells (see Video S3). **B.** Live imaging on a *Tg(krt5:Gal5-ERT, UAS:meGFP)* embryo in which the olfactory orifice (indicated with a white arrowhead) opens through the local loss of cell/cell contacts at the junctions between 4 peridermal cells, highlighted with colors (see Video S4). **C.** Histogram presenting the proportion of live imaged embryos showing local loss of cell/cell contacts, cell division or cell death at the exact location of the orifice and at the time of orifice formation (n = 44 movies from 14 independent experiments). **D.** Graph showing the evolution of the orifice areas as a function of time during the two hours following their opening. Two independent clutches are shown, with n = 5 embryos (clutch 1, black curves) and n = 6 embryos (clutch 2, grey curves). **E.** Pie chart presenting the proportion of live imaged embryos in which the orifice opened at the junction between 3, 4, 5 or more peridermal cells (n = 25 movies in which peridermal cells were labelled with membrane GFP, from 7 independent experiments). The two pictures on the right show additional examples of orifices opening at the junctions between 3 (top) and 4 (bottom) peridermal cells. **F.** The skin epithelium was segmented to compare the area and circularity of skin cells near the forming orifice (blue) with skin cells located further away from the orifice (orange). The measurements were carried out on movies of *Tg(krt5:Gal4-ERT, UAS:meGFP)* embryos, right after the opening (n = 14 movies from 3 independent experiments). The white arrowhead indicates the forming orifice. Error bars: standard deviation. Unpaired, two-tailed t test. **G.** The height of peridermal skin cells was measured using movies performed on *Tg(krt5:Gal4-ERT, UAS:RFP)* or *Tg(krt5:Gal4-ERT, UAS:GFP)* embryos, which were resliced to obtain a transverse view of the epithelium (n = 13 movies from 5 independent experiments). Error bars: standard deviation. Unpaired, two-tailed t test. Scale bars: 20 µm.

In movies where the peridermal cells were labelled with membrane GFP (n = 25 movies from 7 independent experiments), we further noticed that the hole does not form at the junction between 2 cells, but rather at the level of a vertex including 3, 4 or even more cells (Figure 2B, E and Video S4). This prompted us to compare the vertices in the skin epithelium overlying the olfactory placode with those of the skin located further away: the skin epithelium covering the placode was slightly enriched in high fold (4, 5) vertices as compared with the epithelium which surrounds the placode (Figure S1). In addition, the cells lining the future orifice were deformed at the opening onset: they had lost the typical polygonal shape of squamous epithelial cells, they were often smaller and rounder than cells located further away from the forming orifice, and sometimes emitted protrusions towards the cavity (Figure 2F, G and Video S1). These observations indicate that the skin epithelium undergoes a local remodelling above the olfactory placode prior to the opening of the olfactory orifice, characterised by cell shape changes and an enrichment of high fold junctions. The rounding up of the skin cells in this area suggests that these multicellular junctions are different from the high-fold vertices shown to be preferential sites for transient epithelial opening or radial cell intercalation in other systems, where no cell deformations were reported (Isasti-Sanchez et al., 2021; Ventura et al., biorxiv 2021, Collins et al., 2021; Bosveld et al., 2018). These results show that, following a local pre-remodelling of the skin epithelium, the opening of the olfactory orifice occurs through the loss of contacts between skin cells. We then analysed the relationship between the skin and the neurons during the opening of the orifice.

### The olfactory orifice opens above the rosette cavity centre

We first mapped the precise position of the skin opening in relation with the radial organisation of the underlying rosette of placodal neurons. Strikingly, in movies where both the periderm layer and the olfactory placode neurons were visualised, the orifice always opened close to the centre of the placodal rosette, above the rosette cavity in 100% of the cases (n = 17 movies from 5 independent experiments, Figure 3A and Video S5). In other words, the centre of the rosette prefigured the position of the opening. During the few hours following the opening, the size of the orifice in the skin correlated with that of the rosette cavity in fixed samples, with a gradual opening of the skin epithelium above the rosette cavity (Figure 3B-D and Video S2, n = 123 embryos from 3 independent experiments). In addition, the rosette cavity was significantly larger when the skin orifice is open as opposed to closed, even in same-stage embryos (Figure 3E), suggesting that a minimal size of the rosette cavity (∼ 90 µm^2^, Figure 3E) is required for the opening, and/or that the cavity area increases during the opening. Taken together, our observations reveal a close spatio-temporal coordination between the widening of the rosette cavity and the opening of the skin orifice, which suggests an instructive role for the placodal rosette in the opening of the skin epithelium. We thus analysed the functional interactions between the rosette and the overlying peridermal skin cells.

**Figure 3.**
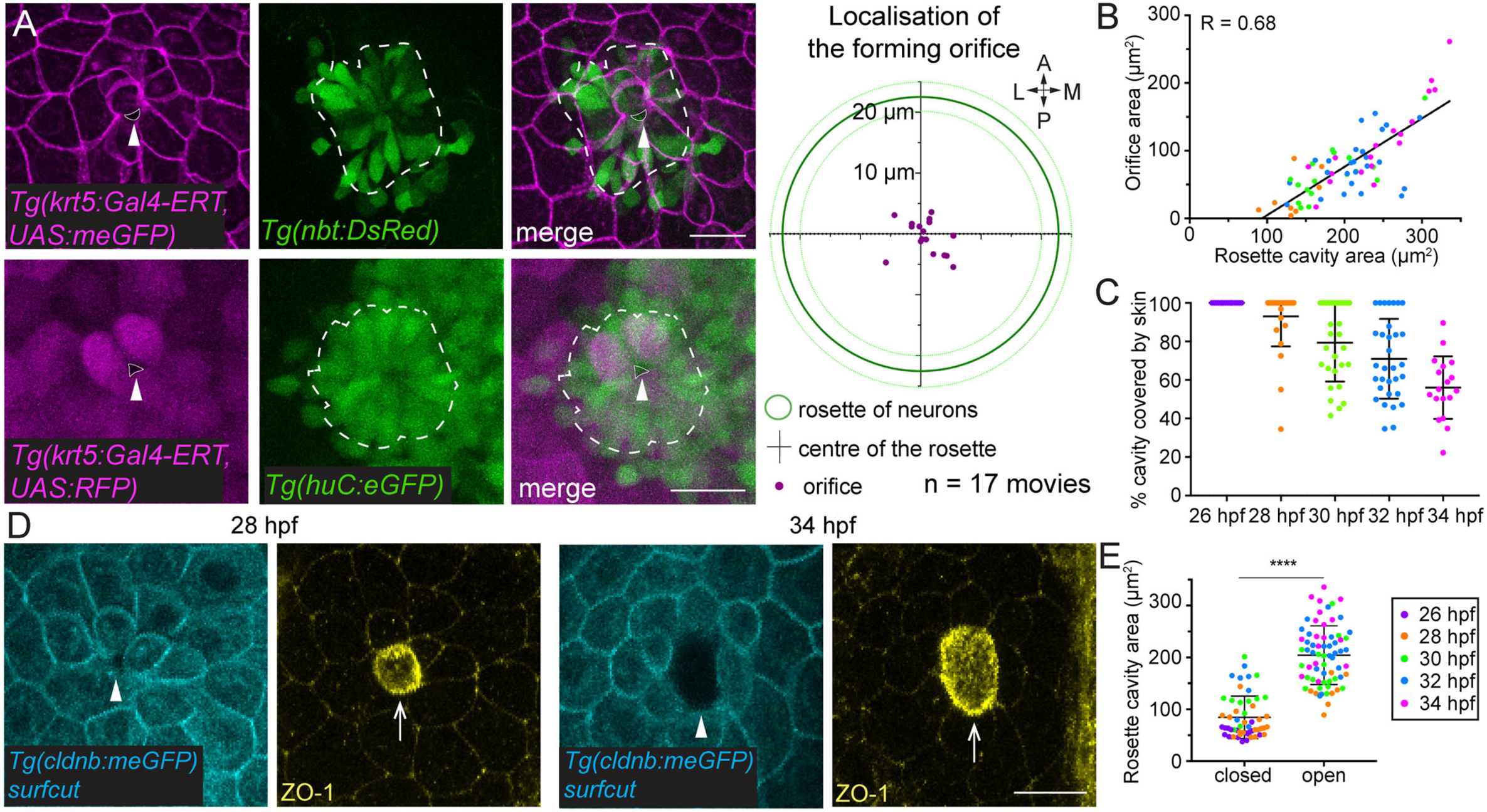
**Interaction between the skin cells and the olfactory neurons during the opening of the orifice**. All images shown in this Figure are dorsal views, oriented anterior to the top, and lateral on the left. **A.** Images extracted from movies performed on embryos in which peridermal cells (magenta) and rosette neurons (green) are labelled with two distinct colors (the transgenic lines employed are indicated on the pictures). The extracted images correspond to the exact time point where the olfactory orifice opens in these embryos, and show that the orifice opens very close to the centre of the underlying rosette structure. The white arrowheads indicate the forming orifices. The contours of the rosettes and of the orifices are shown with dashed and dotted lines, respectively. The right panel shows a graph presenting the position of the orifice opening, in relation with the position of the centre of the underlying rosette (n = 17 movies from 5 independent experiments). The green continuous circle indicates the mean size of the rosette, the dotted circles represent the standard deviation. **B.** Graph showing the correlation between the area of the skin orifice and that of the underlying rosette cavity, analysed on *Tg(cldnb:meGFP)* embryos fixed at various stages and immunostained for GFP and ZO-1, as shown in D. The colors correspond to the stages of development indicated in the legend of the graph in E (n = 123 embryos, 3 independent experiments). **C.** Graph showing the percentage of the rosette cavity area covered by the skin (see Methods), at different developmental stages. The colors correspond to the stages of development indicated in the legend of the graph in E. **D.** Examples of images of 28 and 34 hpf *Tg(cldnb:meGFP)* embryos immunostained for GFP (cyan) and for the tight junction protein ZO-1 (yellow), used to generate the graphs shown in B, C, E. The image J plugin SurfCut (Erguvan et al., 2019) was used to extract the periderm layer above the placode and analyse the olfactory orifice (see Methods). The ZO-1 immunostaining was used to analyse the rosette cavity. The arrowheads point to the open orifices, and the arrows show the underlying rosette cavities. **E.** Graph comparing the size of rosette cavities in placodes with closed versus open skin orifices. Error bars: standard deviation. Unpaired, two-tailed t test. Scale bars: 20 µm.

### The placodal rosette instructs the opening of the olfactory orifice in the skin

To perturb the organisation of the rosette, we manipulated the early coalescence of the olfactory placodes, during which morphogenetic cell movements transform two elongated placodal cell fields lying on each side of the brain into spherical neuronal clusters (Breau et al., 2017; Aguillon et al., 2020; Monnot et al., 2022). We reasoned that perturbing the morphogenesis of the olfactory placode at these early stages (12-22 hpf) would likely result in abnormal rosette formation at our stages of interest (26-34 hpf). To do so, we used a validated morpholino targeting Cxcl12a (Sdf1a) (David et al., 2002), a chemotactic cue produced by the brain and acting on the Cxcr4b-expressing olfactory placode cells to control their anteroposterior convergence movements during placode coalescence (Miyasaka et al. 2007; Aguillon et al., 2020). The knockdown of the Cxcl12a cue should not affect the skin directly, since the Cxcr4b receptor is expressed in the olfactory placode cells, but not in the skin. In this experiment, we mostly used the shape and size of the rosette cavity, labelled with ZO-1 immunostaining, as a readout for the organisation of the rosette (Figure S2 shows images illustrating the correspondence between the organisation of the rosette and of its cavity). At 34 hpf, when in embryos injected with the control morpholino most of the rosette cavities exhibited sizes and shapes that are expected at this stage (Figure 4A1, 55 control embryos), 34% of the Cxcl12a morphants showed defects in their rosette cavities (29 out of 86 embryos, 3 independent experiments), confirming that the organisation of the rosette can be affected in this condition. The phenotypes in Cxcl12a morphants included a few elongated rosette cavities (aspect ratio (AR) > 2, n = 7 embryos), and the presence of two - or three in one sample - distinct cavities in one placode, exhibiting similar or different sizes (n = 22 embryos) (Figure 4A2-A5 and Figure S2). For the following analyses, we selected these abnormal rosette cavity organisations and studied the effect on the opening of the olfactory orifice in the skin. Above the cavities of normal size observed in Cxcl12a morphants (> 90 µm^2^), whether elongated or found in double/multiple cavity situations, the skin epithelium was open, and the localisation and size of the olfactory orifices precisely matched with those of the underlying rosette cavities, as observed in controls (Figure 4A1-C1, A3-C3, A4-C4, and D-F). The matching between the two structures suggests that the communication between the rosette cavity and the skin is conserved when the rosette organisation is affected in Cxcl12a morphants, as expected if the rosette cavity is essential for the opening of the orifice. In situations of double/multiple cavities, a lot of small cavities (< 90 µm^2^) were observed, above which the skin was not open (Figure 4A3-C3, A4-C4 and Figure 4G, orange crosses), whereas almost all the olfactory orifices were open in the control embryos (except in 2 out of 55 embryos, above small cavities, see Figure 4G). This observation reinforces the idea that the orifice opening can only initiate when the skin epithelium covers a sufficiently large rosette cavity. Strikingly, in one sample, the presence of two large rosette cavities was associated with the formation of two distinct holes in the overlying skin epithelium (Figure 4A5-C5 and Video S6), showing that an additional and wide-enough rosette cavity is sufficient to initiate the formation of an orifice in the overlying skin. Altogether, these results support the idea that the placodal rosette instructs the skin opening process.

**Figure 4.**
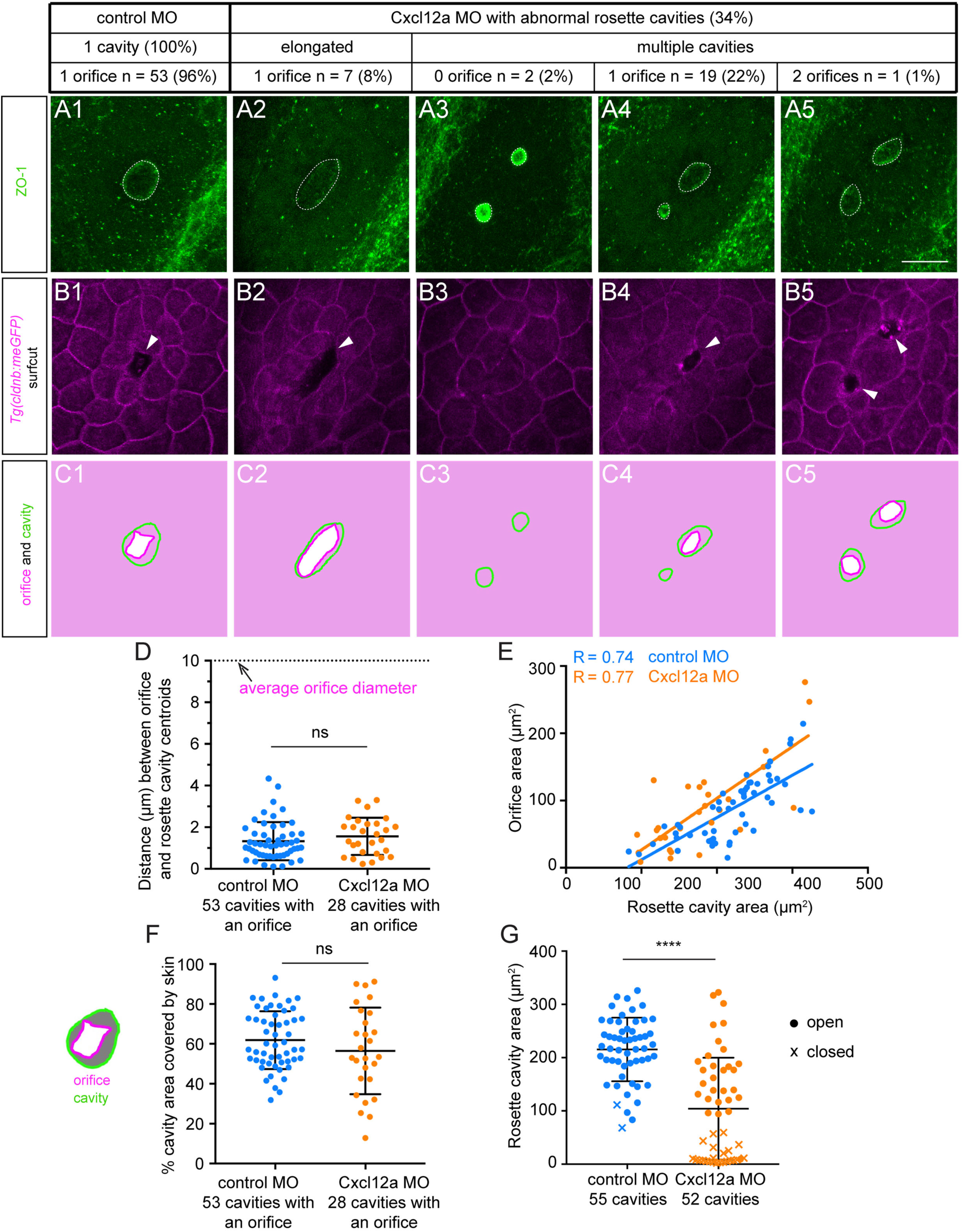
The placodal rosette instructs the opening of the olfactory orifice in the skin. All images shown in this Figure are dorsal views oriented anterior to the top, and lateral on the left. **A, B.** Images of representative 34 hpf *Tg(cldnb:meGFP)* embryos injected with standard control or Cxcl12a morpholinos, immunostained for the tight junction protein ZO-1 (in green) to visualise the rosette cavities in A, and for GFP (in magenta) to visualise the olfactory orifice in B (the most superficial layer has been extracted with the SurfCut plugin, see Methods). The rosette cavities are surrounded by dotted lines. The orifices in the overlying skin are indicated with arrowheads. Cxcl12a morpholino-injected embryos exhibit elongated rosette cavities (AR > 2), as well as double cavities of similar or different sizes. Scale bar: 20 µm. **C.** The contours of the rosette cavities and of the olfactory orifices are shown in green and magenta, respectively. The skin layer covering the placode is colored in light pink. **D.** Graph presenting the distance between the centroids of the rosette cavities and the centroids of the olfactory orifices in 34 hpf control embryos (blue dots) and Cxcl12a morphants (orange dots). Error bars: standard deviation. Unpaired, two-tailed t test. **E.** Graph showing the correlation between the area of the skin orifice and that of the underlying rosette cavity in control embryos and Cxcl12a morphants at 34 hpf. **F.** Graph showing the proportion of the rosette cavity area covered by the skin (grey area in the schematic view, see Methods), in control embryos and Cxcl12a morphants 34 hpf. Error bars: standard deviation. Unpaired, two-tailed t test. **G.** Graph showing the size of the rosette cavities in control embryos and Cxcl12a morphants 34 hpf, with dots indicating rosette cavities above which the orifice is open, and crosses indicating rosette cavities above which the skin is closed. Error bars: standard deviation. Unpaired, two-tailed t test.

### Actomyosin distribution and dynamics suggest a role for mechanical forces in the opening of the olfactory orifice

Our findings suggest so far that the communication from the olfactory placode to the skin, whether chemical and/or mechanical, is crucial for the opening of the olfactory orifice. To get insights into the mechanical cues at play, we first analysed the subcellular localisation of actomyosin, the main force-producing machinery in cells and tissues, right before and during the opening, on 27-29 hpf embryos. We visualised F-actin with injection of utrophin-GFP mRNA or with fluorescent phalloidin, and myosin II with the *Tg(actb2:myl12.1-eGFP)* reporter line, in which a GFP-tagged myosin II regulatory light chain is expressed under the control of the ubiquitous *actb2* promoter (Maître et al., 2012). In peridermal skin cells, actomyosin localised in the apical microridges (Raman et al., 2016) and at the cell/cell junctions between peridermal epithelial cells (Figure 5A-C’ and Video S7). In the placodal cells forming the rosette, two pools of actomyosin were detected. Actomyosin components were first observed to strongly accumulate at the apical surface of the dendrites, forming a multicellular cup-shaped network all around the hemispheric rosette cavity (Figure 5A-C’ and Video S7). In addition, we detected radial actomyosin cables in the dendrites, that most of the time appeared to contact the actomyosin cup lining the rosette cavity. Their length was variable, spanning part of the dendrites in rosette neurons and in some cases extending further down to the cell bodies (Figure 5A-C’ and Video S7).

**Figure 5.**
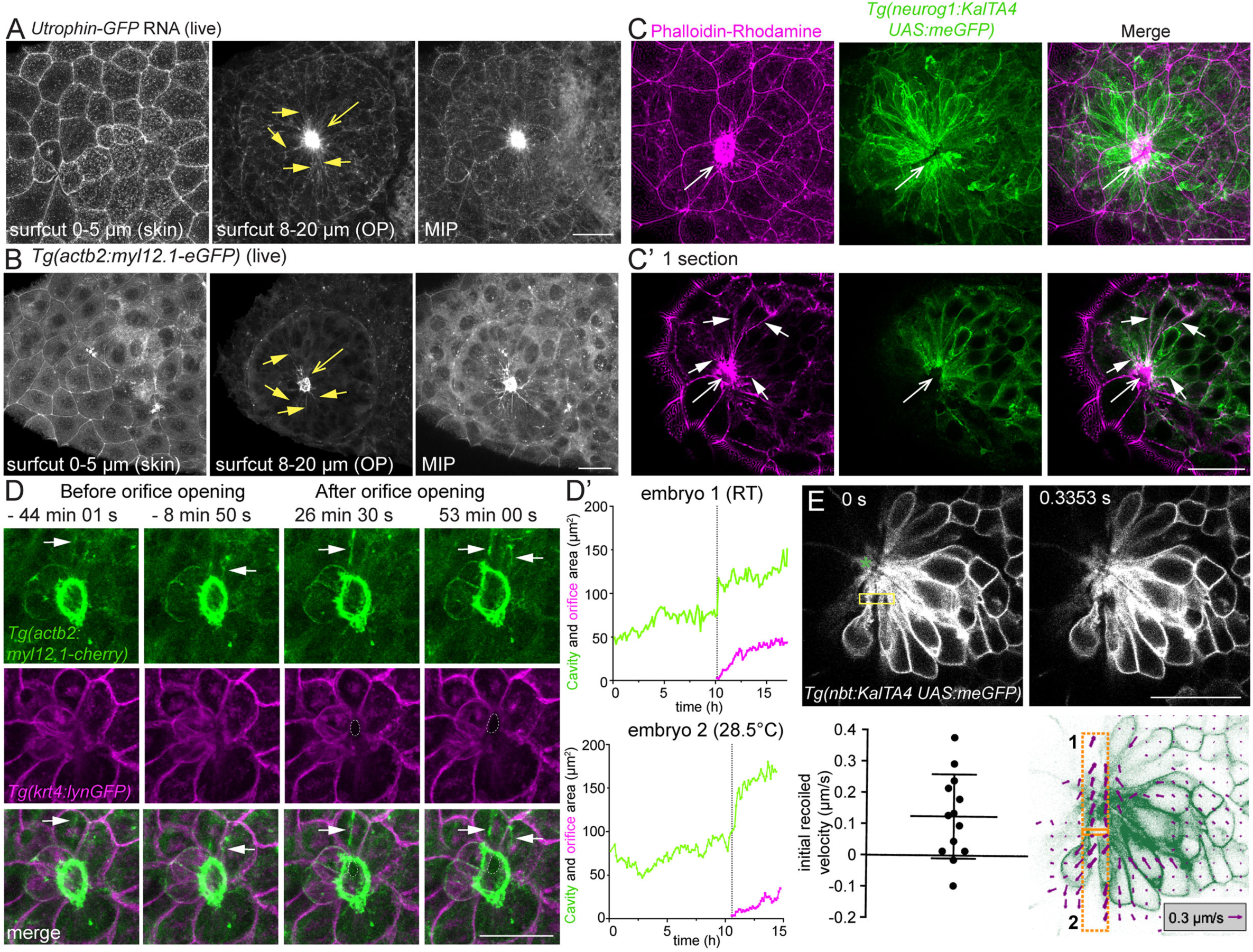
Actomyosin distribution and dynamics in skin cells and rosette neurons. All images shown in this Figure are dorsal views oriented anterior to the top, and lateral on the left. **A, B.** Images of 28 hpf embryos injected with Utrophin-GFP mRNA to visualise F-actin (A) or expressing the *Tg(actb2:myl12.1-eGFP)* transgene to visualise myosin II (B). For each staining, the left picture shows the extraction of the skin peridermal layer using the SurfCut plugin (0-5 µm from the edge of the embryo, see Methods), the middle picture shows the extraction of the dorsal surface of the olfactory placode, where the rosette and its cavity are located (8-20 µm of SurfCut extraction from the edge of the embryo), and the right picture shows the total z maximum intensity projection (MIP). The arrow with an empty head indicates the accumulation of actin and myosin II at the apical tips of the dendrites in the rosette, forming an actomyosin cup structure lining the rosette cavity. The other arrows with filled heads show examples of radial actomyosin cables observed in the placode close to the apical actomyosin cup. **C, C’.** Image of a 28 hpf *Tg(neurog1:KalTA4, UAS:meGFP)* (green) stained with Rhodamine-Phalloidin to label F-actin (magenta). The top panel (C) shows the total z maximum intensity projection and the bottom panel (C’) shows 1 z-section at the level of the placodal rosette. The arrow with an empty head indicates the actomyosin cup. The other arrows with filled heads show examples of radial actomyosin cables seen in the placodal cells forming the rosette. See also Video S7. **D, D’.** Images extracted from a movie performed on an embryo carrying the *Tg(krt4:lynGFP)* and *Tg(actb2:myl12.1-mCherry)* transgenes, allowing the visualisation of skin cell membranes (magenta), and myosin II distribution (green), respectively. Note that the actomyosin cup lining the rosette cavity expands during and after the opening of the olfactory orifice (the orifice is surrounded by dotted lines). The graphs in D’ show the evolution of the actomyosin cup area (green curve) and of the olfactory orifice area (magenta curve) as a function of time in two embryos. The images shown in D correspond to embryos 1 and 2. One experiment was performed at Room Temperature (RT) and the other at 28.5°C. The vertical dotted line indicates the time point at which the olfactory orifice opened in the embryo. The opening of the olfactory orifice is concomitant with a sudden increase in the area of the actomyosin cup structure lining the rosette cavity. The results obtained for 5 additional embryos are shown in Figure S3. **E.** Laser ablations were performed in small areas located in the sub-apical regions of the rosette dendrites at 28-30 hpf, meaning right before or during the olfactory orifice opening. The dendrites and cell bodies of the rosettes neurons were visualised with the *Tg (nbt:KalTA4; UAS:meGFP)* line. The left image illustrates the position of the region of ablation (yellow), just before the ablation. The position of the rosette cavity is indicated with a green asterisk. 0s corresponds to the last time point before the ablation. The right image shows the tissue velocity field overimposed (magenta arrows) in 20 µm-long rectangles on both sides of the ablated region. The recoil velocity was obtained with the following formula: V_recoil_ = <V^┴^>_2_ - <V^┴^>_1_ where V^┴^ indicates the average outward velocity component orthogonal to the cut orientation, and < > denotes the average over the adjacent rectangle (1 or 2). We arbitrarily oriented the orthogonal axis towards the outside of the rosette. The graph shows the initial recoil velocity, used as a proxy for dendritic tension before the cut, following the ablations (n = 13 placodes, 1 ablation/placode, data obtained from 4 independent experiments). Error bar: standard deviation. Scale bars: 20 µm.

To capture the dynamics of actomyosin before and during the orifice opening, we performed live imaging using the *Tg(actb2:myl12.1-eGFP)* and *Tg(actb2:myl12.1-mCherry)* reporter lines (Maître et al., 2012). In a first set of experiments, we took advantage of a GFP-targeted Crispr/Cas9 approach to generate mosaic *myl12.1-eGFP* expression in the ubiquitous *Tg(actb2:myl12.1-eGFP)* line, which allowed us to better appreciate the dynamics of myosin II in isolated single cells or small groups of cells. High spatiotemporal resolution imaging of mosaic *myl12.1-eGFP* before the opening (28 hpf) highlighted differences in the two pools of myosin II in the placodal neurons: an overall static myosin II pool located at the apical tip of the dendrites (the actomyosin cup), likely representing the actomyosin cortex coupled with the apical adhesion belts, and the radial actomyosin cables which appeared to be more dynamic structures, moving, changing size and assembling/disassembling overtime (Video S8). We next performed live imaging of myosin II and skin cell behaviours during the opening of the orifice. The *Tg(krt5:Gal4-ERT, UAS:RFP*) line was used to visualise skin cells, and since the actomyosin cup surrounds the rosette cavity, the localisation of myl12.1-eGFP in the cup was used as a readout for the cavity. We observed an initial phase of slow growth of the rosette cavity, followed by a sudden expansion of the cavity. Strikingly, this sudden widening was concomitant with the opening of the olfactory orifice in the overlying skin (n = 7 movies) (Figure 5D-D’, Figure S3 and Video S9). In all movies, the few cables assembling during the skin opening seemed to pull radially on the actomyosin cup structure lining the rosette cavity, thus potentially contributing to its expansion (Figure 5D and Video S9).

To further test the hypothesis of radial pulling forces exerted by the rosette neurons, we carried out laser ablation to analyse whether their dendrites are under mechanical tension during the opening of the orifice, at 28 hpf (n = 13 analysed placodes from 4 independent experiments, 1 cut/placode). This approach was previously used in zebrafish to probe the tension of cellular extensions in other systems (MacDonald et al., 2015; Pulgar et al., 2021). In a large majority of the cases, we observed immediate recoil after severing the sub-apical region of the rosette dendrites (Figure 5E), suggesting that they bear mechanical tension and supporting the idea that they could exert active, actomyosin-mediated pulling forces on the overlying skin cells.

### Actomyosin contractility in placode neurons is necessary to open the olfactory orifice

To further investigate the role of actomyosin, we carried out a treatment with blebbistatin, an inhibitor of non-muscle myosin II activity, during the formation of the olfactory orifice, from 26 to 34 hpf. This treatment completely prevented the opening of the orifice in all embryos (Figure 6A, A’, n = 16 embryos treated with DMSO and n = 18 embryos treated with blebbistatin, 3 independent experiments), demonstrating that actomyosin activity is essential for the skin epithelium to open. Of note, the size of the rosette cavities at 34 hpf was smaller than in controls upon blebbistatin treatment (Figure 6A’), but above the threshold required to have the competence to open the olfactory orifice (90 µm^2^, see Figure 3E). This suggests that actomyosin contractility contributes to the expansion of the rosette cavity. To test this hypothesis, we performed live imaging in blebbistatin-treated embryos. Here again we used the localisation of myl12.1-eGFP in the actomyosin cup as a proxy for the rosette cavity, and the *Tg(krt5:Gal4-ERT, UAS:RFP*) line to visualise skin cell behaviours (n = 6 movies). This live imaging experiment first confirmed the absence of skin opening upon blebbistatin treatment. In all embryos, the initial, slow phase of rosette cavity growth was overall maintained, but no sudden and sharp expansion was detected (Figure 6B and Figure S4). These results show that the first phase of progressive growth of the rosette cavity is actomyosin-independent. The lack of a sudden widening of the rosette cavity in blebbistatin-treated embryos could be due to a direct requirement of actomyosin in this step of the cavity expansion, or be a secondary consequence of the absence of hole opening in the skin.

**Figure 6.**
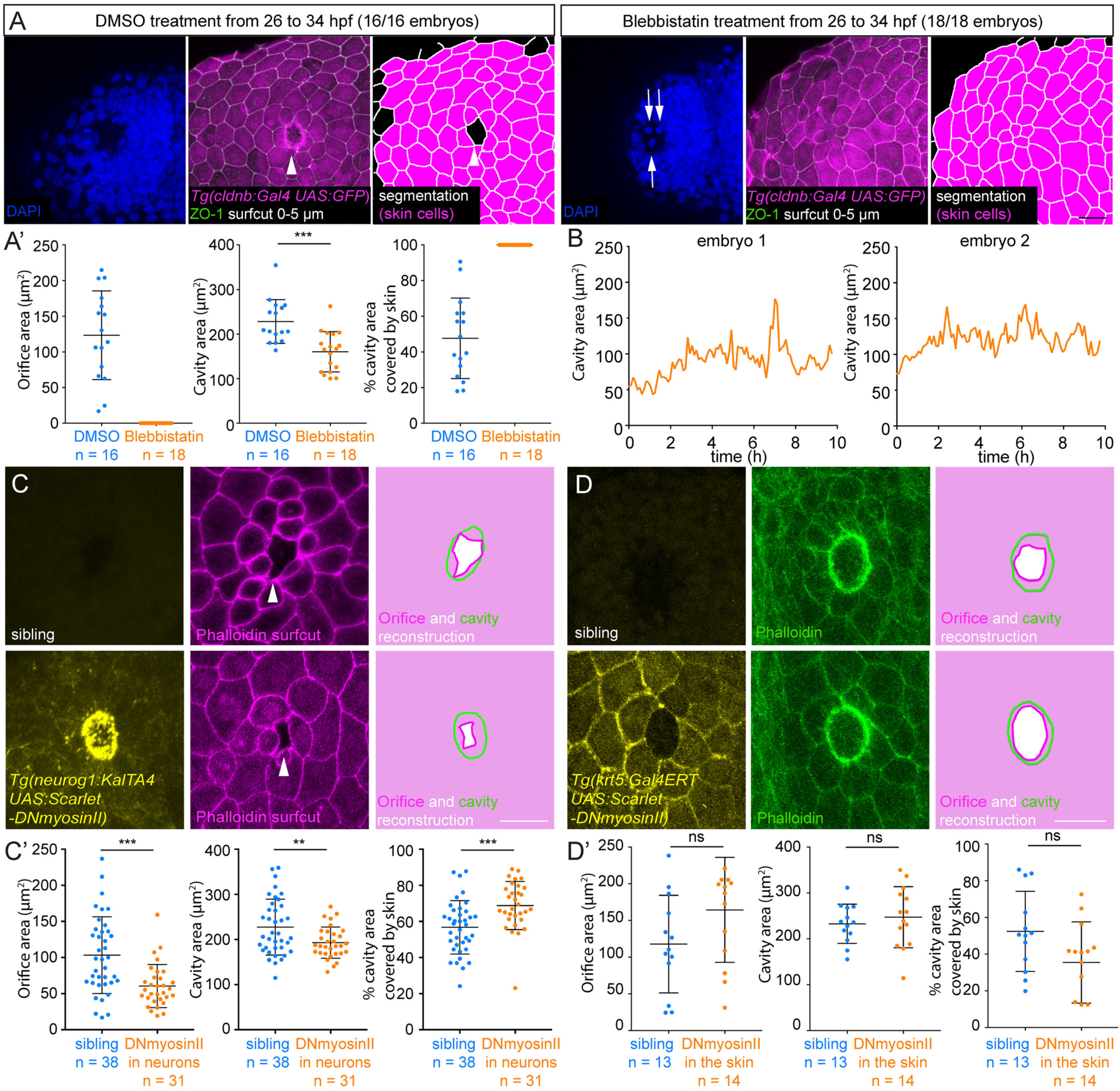
Actomyosin activity is required in the placode neurons to open the olfactory orifice. A. Representative images of 34 hpf *Tg(cldnb:Gal4, UAS:GFP)* embryos treated with DMSO (left panel) or blebbistatin (right panel) from 26 hpf to 34 hpf. In each panel, the left pictures show the nuclear DAPI staining (blue), the middle pictures show the GFP-expressing skin cells (magenta) labelled with ZO-1 (green) (the skin layer was extracted with the SurfCut plugin, 0-5 µm from the edge of the embryo), and the right panels show the segmented contours of skin cells and of the olfactory orifices (using the ZO-1 labelling). While all the orifices were open in DMSO control embryos (n = 16), no orifice was found in the blebbistatin-treated embryos (n = 18), 3 independent experiments. **A’**. Graphs showing the area of the olfactory orifices, the area of the rosette cavities, and the proportion of the rosette cavity covered by the skin in blebbistatin- and DMSO-treated embryos at 34 hpf. Unpaired, two-tailed t-test. Error bar: standard deviation. **B.** Movies were performed on *Tg(krt5:Gal4-ERT, UAS:RFP*)*; Tg(actb2:myl12.1-eGFP)* embryos treated with blebbistatin from 26 hpf, and the area of the actomyosin cup was analysed as a function of time. Note that in these embryos, the olfactory orifice does not open. The quantification for the 4 other embryos is shown in Figure S4. **C**. Images of embryos expressing mScarlet-DN-MYL9, a dominant negative form of myosin II (DN myosin II), specifically in placodal neurons, and control siblings at 34 hpf. The expression of mScarlet-DN-MYL9 is shown in yellow, and the phalloidin staining (SurfCut extraction of the skin, see Methods) in magenta. The reconstructions of the rosette cavities and the olfactory orifices are depicted in the right column. Scale bars: 20 µm. **C’**. Quantification of the olfactory orifice areas and rosette cavity areas in both conditions, as well as the proportion of the cavity area covered by skin cells. For the cavity area, an unpaired, two-tailed t-test was done. For the orifice area and the percent of cavity area covered by skin, unpaired Mann-Whitney tests were performed. Error bar: standard deviation. **D.** Images of embryos expressing mScarlet-DN-MYL9 specifically in peridermal skin cells, and control siblings at 34 hpf. The expression of mScarlet-DN-MYL9 is shown in yellow, and the phalloidin staining in green. The reconstructions of the rosette cavities and the olfactory orifices are depicted in the right column. Scale bars: 20 µm. **D’.** Quantification of the olfactory orifice areas and rosette cavity areas in both conditions, as well as the proportion of the cavity area covered by skin cells. Unpaired, two-tailed t-test. Error bar: standard deviation.

To further investigate this aspect, we manipulated actomyosin activity specifically in the olfactory placode or in the skin by using the Gal4/UAS system to express a dominant-negative form of myosin II (mScarlet-DN-MYL9) known to reduce actomyosin contractility in other contexts (Beach et al., 2011; Priya et al., 2020). We generated a *UAS:mScarlet-DN-MYL9* construct, performed Tol2-mediated transgenesis, and used three independent *Tg(UAS:mScarlet-DN-MYL9)* F0 founders transmitting the transgene to the progeny. To achieve neuron-specific expression, we produced a *Tg(neurog1:KalTA4)* line, expressing KalTA4 (Distel et al., 2009) in the early-born neurons of the olfactory placode. In 34 hpf embryos expressing mScarlet-DN-MYL9 specifically in the placodal neurons, the size of the olfactory orifice was significantly reduced as compared with control siblings (Figure 6C, C’, n = 31 embryos expressing mScarlet-DN-MYL9 in neurons, and n = 38 control siblings, 3 independent founders, 2 independent experiments). Consistently, the percentage of the cavity area covered by the skin monolayer was increased (Figure 6C’). As observed in blebbistatin-treated embryos, the size of the rosette cavities was smaller, but above the threshold required to be competent to open the skin (90 µm^2^, see Figure 3E). These results demonstrate that actomyosin in the olfactory placode neurons is necessary for a normal, full opening of the overlying skin epithelium, and suggest that it mediates the sudden expansion of the rosette cavity accompanying the opening of the orifice. By contrast, expressing mScarlet-DN-MYL9 specifically in the periderm with the *Tg(krt5:Gal4-ERT)* driver had no effect on the opening of the orifice nor the size of the rosette cavity (Figure 6D, D’, n = 14 embryos expressing mScarlet-DN-MYL9 in the periderm, and n = 13 control siblings, 2 independent experiments), suggesting that actomyosin activity is not intrinsically required in the skin for the formation of the olfactory orifice. Collectively, these tissue-specific perturbations show that actomyosin contractility is required specifically in placodal neurons to fully open the olfactory orifice in the skin, in agreement with an implication of out-of-plane, extrinsic pulling forces in the epithelial opening process.

## Discussion

While the closure of epithelial gaps during development and wounding has been extensively studied (reviewed in Begnaud et al., 2016; Zulueta-Coarasa and Fernandez-Gonzalez, 2017), the formation of breaks or holes in epithelial tissues remains poorly understood and has recently received a growing attention in the field of epithelial mechanics (Harris et al., 2012, Casares et al., 2015; Sonam et al., biorxiv; Dumortier et al., 2019; Proag et al., 2019; Fouchard et al., 2020; Isasti-Sanchez et al., 2021; Prakash et al., 2021; Row et al., 2021, Bonfanti et al., 2022). Our study shows that the opening of the olfactory orifice in the skin epithelium is instructed by the underlying rosette of neurons located in the dorsal olfactory placode. Our results are compatible with a model in which the rosette neurons contact the overlying skin monolayer through the apical tip of their dendrites, and pull radially on the skin to locally break the epithelium. This highlights a novel mechanism for the physiological opening of an epithelial barrier (Figure 7).

**Figure 7.**
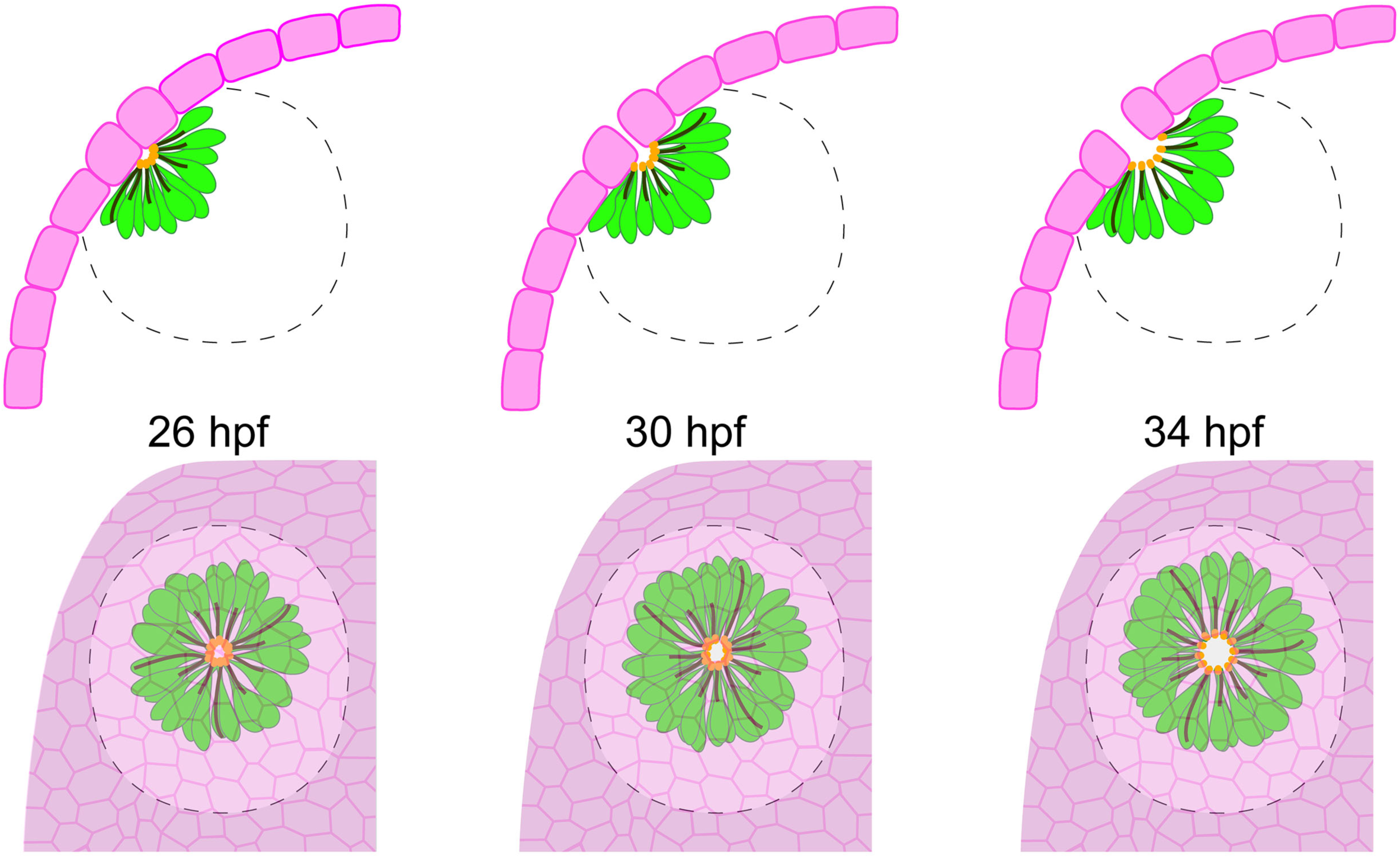
Mechanisms driving the opening of the olfactory orifice. This schematic view depicts our findings and the proposed model. The three drawings represent the 26, 30 and 34 hpf stages. Before the opening onset, at 26 hpf, the olfactory placode neurons (in green) are assembled into a rosette structure which is juxtaposed to the overlying monolayer of peridermal skin cells (in magenta). Actomyosin accumulates at the apical tips of the rosette dendrites, forming an actomyosin cup structure lining the rosette cavity in the rosette centre (orange). Dynamic, radial cables of various sizes (black) are also observed in the olfactory placode neurons. From around 30 hpf, some of these cables appear to pull on and deform the actomyosin cup structure as well as the entire rosette cavity, thus mediating its sudden expansion. The sudden widening of the rosette cavity induces, through physical contact, a local loss of cell/cell adhesion in the upper skin layer which marks the onset of the opening of the orifice and exposes the sensory neurons to the outside. These findings unravel a previously uncharacterised biomechanical process driving the opening of an epithelial barrier, whereby extrinsic forces exerted by an adjacent cell population tear epithelial cells to trigger hole formation.

Similar mechanisms could act widely in sensory organ morphogenesis. As mentioned earlier, the development of sensory organs such as the taste buds and the lateral line neuromasts requires the formation of a pore in epithelial layers, allowing the sensory cells to access external cues (Hansen et al., 2002; Gleason et al., 2009; Barlow, 2015; Dow et al., 2018; Hino et al., 2022). This may involve the rosette structures formed by sensory cells and support cells in these two types of sensory organs, as well as mechanical or chemical communication with the skin or the tongue epithelia, but the underlying mechanisms remain to be investigated. The amphid organ in *C. elegans* is another sensory organ which develops through the physical interaction of the skin with a multicellular rosette of neurons and glia (Fan et al., 2019). Here, the forces have been described to be exerted by the skin on the neuronal rosette: the anterior movement of the epidermis pulls on the dendrite tips of the neurons forming the rosette, thus extending them and dragging them to the future location of the nose. This is followed by the posteriorward movement of the cell bodies further extending the dendrites of the amphid neurons (Heiman and Shaham, 2009; Fan et al., 2019). The molecular mechanisms of attachment between the two cell types have been well studied in this context (Heiman and Shaham, 2009; Fan et al., 2019). However, how the epidermis opens to expose the amphid neurons to the environment remains unclear, in particular whether the opening involves the amphid dendrites themselves or the surrounding glial cells (Low et al., 2019). A common feature in the morphogenesis of sensory organs is thus the formation of stable, multicellular rosettes of sensory cells which are attached to the overlying epithelial cells. The radial organisation of their cells could facilitate interactions with the adjacent epithelial layers to promote the opening of the gaps for the detection of external sensory stimuli.

The formation of gaps in epithelial tissues also occurs during cell intercalation events, such as the extravasation of immune cells through the endothelial barrier (Escribano et al., 2019), or radial cell intercalation within the *Xenopus* embryo epidermis, in which multiciliated cells integrate into the superficial skin through the local loss of homotypic cell-cell junctions (Stubbs et al., 2006; Sedzinski et al., 2016; Chuyen et al., 2021; Collins et al., 2021). In the *Xenopus* epidermis, the multiciliated cells induce the pre-remodelling of the junctions in the superficial epithelial layer to promote the formation of high order junctions, which prime the epithelium for their further integration (Ventura et al., biorxiv). Interestingly, we found that the olfactory orifice preferentially opens at the junction between 3 or 4 peridermal skin cells. However, since these cells are round and lost their typical epithelial morphology, the high fold junctions we observe are likely endowed with structural and mechanical properties that are distinct from the canonical high fold vertices described in the *Xenopus* epidermis or in other epithelial systems (Bosveld et al., 2018; Collins et al., 2021; Ventura et al., biorxiv). We hypothesise that the deformation and rounding up of peridermal cells prior to the opening represents a permissive state of the skin epithelium facilitating the initial encounter with the underlying neurons and the local rupture of cell/cell contacts in the skin. The potential influence of the placodal rosette on this periderm pre-remodelling will need to be elucidated. In addition, as it has been shown that holes spontaneously form in epithelial regions with low cell/ECM adhesion *in vitro* (Sonam et al., biorxiv), it will be interesting to investigate whether the absence of basement membrane and basal cells above the olfactory placode promotes hole opening in the peridermal skin cells.

Whereas the *Xenopus* multiciliated intercalating cells send filopodia towards the intercellular junctions of the surface epithelium (Ventura et al., biorxiv), we did not detect any sign of protrusive activity from the placodal neurons directed towards the skin. In the placodal rosette, the tips of the dendrites are tightly connected to each other through apical adhesion belts lining the rosette cavity, as illustrated by ZO-1 localisation, which likely prevents the neurons from sending protrusions and pulling individually on the skin epithelium, as do isolated multiciliated cells in the *Xenopus* epidermis. In such a collective rosette organisation, following the initial contact between the skin and the neurons, only a change in the size, shape, or a movement of the rosette and its internal cavity could pull the skin cells above. We found that the opening of the orifice is associated with a sudden dilation of the actomyosin cup lining the rosette cavity. During this expansion, the radial cables present in the placode neurons appear to pull on and expand the apical actomyosin network and likely the entire rosette cavity, thus inducing, through physical coupling, the opening in the overlying skin epithelium. We propose that this sequence of events represents the force-producing machinery pulling radially on the skin cells and triggering hole formation in the epithelium (Figure 7). This model fits with the notion of a minimal cavity size required for the formation of an olfactory orifice, which may represent a threshold of mechanical forces to reach for the opening to initiate.

Other examples of epithelial ruptures in response to force loading have been described in the recent literature, including holes in epithelial tissues of *Trichoplax adhaerens* induced by its own motility behaviour (Prakash et al., 2021), or holes spontaneously forming in epithelial monolayers *in vitro* (Sonam et al., biorxiv). In these examples, the fractures result from an autonomous tensile stress within the epithelial tissue itself, building up upon local contractions (Armon et al., 2018; Prakash et al., 2021) or low adhesion to the substrate (Sonam et al., biorxiv). Our findings unravel a distinct *in vivo* mechanism for epithelial disruption whereby out-of-plane forces applied by an adjacent cell population tear epithelial cells to trigger hole formation.

## Methods

### Fish lines

Wild-type and transgenic zebrafish embryos were obtained by natural spawning. In the text, the developmental times in hpf indicate hours post-fertilisation at 28.5°C. We used the following lines (simplified names are used in the figures and their legends): *Tg(krt5:Gal4-ERT2-VP16, myl7:CFP)^b1234^* (Akerberg et al., 2014) and *Tg(krt4:LY-eGFP)^sq18^*(a gift from Tom Carney, Singapore) (Lee et al., 2014) to label and target peridermal cells, *Tg(-4.0cldnb:GalTA4, cry:RFP)^nim11^*(Breau et al., 2013) and *Tg(-8.0cldnb:LY-EGFP)^zf106^* (Haas and Gilmour, 2006) to label both olfactory placode and peridermal cells, *Tg(Xla.Tubb:DsRed)^hkz018t^*(Peri and Nüsslein-Volhard 2008); *Tg(-2.0ompb:gapYFP)^rw032^* (a gift from Nobuhiko Miyasaka, RIKEN Institute, National Bioresource Project of Japan) (Sato et al., 2005), *Tg(-8.4neurog1:GFP)^sb1^* (Blader et al., 2003), *Tg(elavl3:GFP)^knu3^*(Park et al., 2000) and the Tg(*Xla.Tubb:KalTA4)* line *- Tg(nbt:KalTA4)* in the text - (a gift from David Lyons) to label and target olfactory placode neurons, *Tg(14XUAS:MA-GFP)^ue102^* (Münzel et al., 2012), *Tg(14XUAS:mRFP,Xla.Cryg:GFP)^tpl2^* (Balciuniene and Balciunas, 2013) and *Tg(UAS:GFP)* (a gift from David Wilkinson) to label the membrane or cytoplasm of Gal4-expressing cells, and *Tg(actb2:myl12.1-eGFP)^e2212^*and *Tg(actb2:myl12.1-cherry)^e1954^* (Maître et al., 2012) as reporter lines to visualise myosin II. The *Tg(-8.4neurog1:KalTA4)* and *Tg(UAS:mScarlet-DN-MYL9)* lines were generated by Tol2-mediated transgenesis (see plasmid construction for details about the cloning steps). All our experiments were made in agreement with the European Directive 210/63/EU on the protection of animals used for scientific purposes, and the French application decree “Décret 2013-118”. The fish facility has been approved by the French “Service for animal protection and health”, with the approval number A-75-05-25.

### Plasmid constructions

The Tol2 -8.4neurog1:KalTA4 plasmid was produced using the gateway system (Kwan et al., 2007), by combining four plasmids: the 5’ entry plasmid with 8.4 kb of the *neurogenin1* promoter (a gift from Patrick Blader), the middle entry KalTA4 (Distel et al., 2009) plasmid (produced in the laboratory of David Lyons), the 3’ entry polyadenylation signal plasmid (Kwan et al., 2007) and the backbone containing the cmcl2:GFP selection cassette (Kwan et al., 2007). The Tol2 UAS:mScarlet-DN-MYL9 plasmid was produced by classical cloning methods: the 5XUAS sequence was amplified by PCR from a medusa plasmid (Distel et al., 2010) using the 5’ CTGACTAGTCTCTAGGGGCTGCAGGTCGG 3’ forward and 5’ CTGGGATCCAGGGCTGCAGAATTCGTGTGG 3’ reverse primers. The UAS amplicon and the Tol2 myl7:mScarlet-DN-MYL9 plasmid (a gift from Rashmi Priya) (Priya et al., 2020) were digested by SpeI and BamHI and ligated using the T4 ligase (Invitrogen).

### Immunostainings

For immunostaining, embryos were fixed in 4% paraformaldehyde (PFA, in PBS), blocked in 3% goat serum and 0.3% triton in PBS for 2h at room temperature or overnight at 4°C and incubated overnight at 4°C with primary and secondary antibodies. The following primary antibodies were used: anti-Tp63 (rabbit, 1/100, Sigma, SAB2701838), anti-Laminin (rabbit, 1/100, L-9393, Sigma), anti-Collagen IV (rabbit, 1/200, Abcam, ab6586), anti-ZO-1 (mouse IgG1, 1/500, 33-910, ThermoFisher), anti-GFP (chicken, 1/200, Aves labs, GFP-1020), anti-DsRed (rabbit, 1/300, Takara, 632496), anti-Parvalbumin (mouse IgG1, 1/200, Sigma, MAB1572). For Phalloidin staining, an overnight incubation was performed at 4°C with Phalloidin-Rhodamin (1/200, ThermoFisher) or Phalloidin-Alexa488 (1/200, ThermoFisher).

### Morpholino, mRNA and Crispr/Cas9 injections

The Cxcl12a ATG blocking morpholino (5’ ATCACTTTGAGATCCATGTTTGCA 3’) (David et al., 2002) and the standard morpholino (5’ CCTCTTACCTCAGTTACAATTTATA 3’) were purchased from Genetools and injected at 0.4 mM in 1-cell stage *Tg(-8.0cldnb:LY-EGFP)* or *Tg(-8.4neurog1:GFP)* transgenic embryos. Injected embryos were incubated at 28.5°C until the shield stage, and placed at 33°C overnight. The next day, they were placed back at 28.5°C overday and fixed at 34 hpf for immunostaining. The embryos injected with the standard morpholino did not show any phenotype as compared with uninjected controls.

Utrophin-GFP mRNA was synthesised from the linearised pCS2-Utrophin-GFP vector (a gift from Marie-Emilie Terret) using the mMessage mMachine SP6 transcription kit (Thermo Fisher Scientific). 30 pg of mRNA were injected in 1-cell stage embryos. Nuclear Cas9 mRNA was synthesised from the linearised pT3TSnCas9n plasmid (Addgene 46757, a gift from Alexis Eschstruth) (Jao et al., 2013) using the mMessage mMachine T3 Transcription Kit (Thermo Fisher Scientific). For the GFP Crispr/Cas9 injection, a mix containing 200 ng/µL GFP RNA guide (Auer et al., 2014) and 150 ng/µL Cas9 mRNA was injected in 1 cell stage *Tg(actb2:myl12.1-eGFP)^e2212^* embryos and at 26 hpf embryos were screen for weak/mosaic GFP expression.

### Drug treatments

For blebbistatin treatment, *Tg(cldnb:Gal4, UAS:GFP)* embryos were collected at 10 am and incubated at 28.5°C up to the shield stage, and then placed at 33°C overnight. The next morning, the embryos were at stage Prim 12-14 at 10 am. They were dechorionated and incubated at 28.5°C with 10 to 50 μM Blebbistatin (Sigma, B0560) or in DMSO only, for 8 hours, to reach the 34 hpf stage (Prim 20-22), and fixed for immunostaining analysis.

To induce the expression of UAS transgenes with the *Tg(krt5:Gal4-ERT)* line, the embryos were treated at 9 hpf with 4-hydroxy-tamoxifen (Sigma, H7904) dissolved in ethanol (final concentration of 8 ng/ μL).

### Image acquisition

For live imaging, embryos were dechorionated manually, tricained and mounted around 26 hpf in 0.5% low melting agarose in 1X E3 medium. Movies were recorded either at 28.5°C on a Leica TCS SP8 MPII upright multiphoton microscope using 25X (numerical aperture (NA) 0.95) or 63X (NA 0.9) water lenses, or at room temperature (24-25°C) on a Zeiss 980 FAST Airyscan II 20X (NA 1.0) water lens. For live imaging in the blebbistatin condition, the drug was added to both the E3 medium and the low melting agarose embedding the embryos. For fixed embryos, immunostained embryos were mounted in 0.5% low melting agarose in PBS and imaged on a Leica TCS SP5 AOBS upright confocal microscope using a 63X (NA 0.9) water lens or on a Zeiss 980 FAST Airyscan with a 20X NA 1.0 water lens.

### Laser ablation of the dendrites

Embryos were dechorionated, tricained and mounted in 0.5% low melting agarose in 1X E3 medium in Ibidi dishes (81158) and imaged using an inverted laser-scanning microscope (LSM 880 NLO, Carl Zeiss) equipped with a 63X oil objective (1.4 DICII PL APO, Zeiss). Rectangular cuts (about 5-10 µm long and 2 µm large) were performed in the sub-apical region of the dendrites, close to the rosette cavity, as depicted in Figure 5E. Ablations were performed using a Ti:Sapphire laser (Mai Tai, DeepSee, Spectra Physics) at 790 nm with <100 fs pulses, using a 80 MHz repetition rate, a 100% power and a number of iterations ranging between 1 and 2. Images were acquired at a frame rate between 0.3 and 1.4 s after ablation. To analyse the relaxation of the dendrites after the cuts, we measured the local flow by particle image velocimetry using the MatPIV toolbox for Matlab (Mathworks, US). The window size was set to 64 pixels (∼10 µm), with an overlap of 0.75. The kinetics of relaxation was assessed between the pre-cut time-point and 1 frame later in two 20 µm-long rectangles on each side of the region of ablation, indicated by the orange rectangles in Figure 5E. The "recoil velocity" was then obtained with the following formula: V_recoil_ = <V^┴^>_2_ - <V^┴^>_1_ where V^┴^ indicates the average outward velocity component orthogonal to the cut line, and < > denotes the average over the rectangle (1 or 2, see Figure 5E). We arbitrarily oriented the orthogonal axis towards the outside of the rosette structure.

### Image analysis

*Olfactory orifice and rosette cavity analysis.* In live or fixed embryos in which peridermal cells were labelled with membrane or cytoplasmic GFP, or with phalloidin staining, the most superficial layer was isolated using the SurfCut plugin (Erguvan et al., 2019). This plugin allows to extract and project, from a 3D sample with a slightly curved geometry (which is the case of the skin overlying the olfactory placode), the fluorescent signal coming exclusively from the sample surface. It also allows to extract the signal of a 3D sample at a given distance from the sample surface, which can be chosen by the user. This feature was used in some cases to extract the most superficial (dorsal) layer of the olfactory placode (Figure 5A, B). After the SurfCut processing, when required, the contours of individual skin cells and of the orifice were semi-automatically segmented with a custom Image J plugin (Sebastian Rosi and Jean-François Gilles), using the peridermal membrane GFP labelling or ZO-1 immunostaining. All the detected contours were checked and manually corrected when required. ZO-1 immunostaining and phalloidin staining were used to analyse the rosette cavities: the area of the rosette cavity was delimited and measured using the freehand selection tool and ROI manager plugin of ImageJ. This step was performed in a blind manner, meaning without visualising the overlying orifice in the skin.

To obtain the graph in Figure 3A, we imaged the neurons and the skin in two distinct colors. At the first time point of orifice opening, the centre of the orifice was reported on a target graph where the reference centre represents the centre of the rosette cavity observed with the apical dendrite tips. Both centres were analysed independently, without knowing where the other centre is located.

*Skin cell measurements.* The ImageJ plugin Slice tool was used to obtain an orthogonal section of the skin at the location of the forming orifice in Figure 2G, to measure the height of the skin cells.

*Percentage of cavity area covered by skin.* The proportion of the cavity covered by the skin was calculated as follows: ((A_cavity_-A_orifice_)/A_cavity_) x 100, where A_cavity_ and A_orifice_ represent the areas in µm^2^ of the rosette cavity and of the olfactory orifice, respectively. This corresponds to the grey area in the schematics shown in Figure 4F.

### Statistical analysis

Graphs show means ± standard deviation, overlayed with all individual data points. The plots were generated with the GraphPad Prism software. For all graphs, we checked for normality of the data distribution before performing parametric unpaired, two tailed t tests. When the data were not normal, a Mann-Whitney test was used. The p values correspond to *p < 0.05, **p < 0.01, ***p < 0.001. No statistical method was used to estimate sample size and no randomisation was performed. Blinding was performed for the analysis of the olfactory orifices and rosette cavities.

## Material availability

All reagents (plasmids and transgenic lines), data and code generated in this study are available upon reasonable request.

## Acknowledgements

We gratefully acknowledge Pierre-Luc Bardet, Estelle Hirsinger, François Robin and Christine Vesque for their insightful comments in reviewing the manuscript, and Sophie Gournet for her help in the artwork. This work was funded by the Agence Nationale pour la Recherche (ANR-17-CE13-0009-01 NEUROMECHANICS), the Centre National pour la Recherche Scientifique (CNRS), Sorbonne Université (including a grant from the i-Bio initiative), and the National Institute of Health NIDCD Grant R01-DC-017989. M. Baraban was supported by a postdoctoral fellowship from the Fondation pour la Recherche Médicale (ARF201809006950). We also thank the imaging platform of the Institut de Biologie Paris-Seine (the facility is supported by CNRS, Sorbonne Université and the Conseil Régional Ile-de-France), the Cell and Tissue Imaging core facility (PICT IBiSA), Institut Curie, member of the French National Research Infrastructure France-BioImaging (ANR10-INBS-04), and the IBPS aquatic platform for fish care.

## Author contributions

MB designed the project. MB and MAB conceived the experiments. MB, CGP, IB, JFG, CL, MC and FT conducted the experiments and/or analysed the data. MB and MAB wrote the article.

## Legends of the Supplementary Data

**Figure S1.**
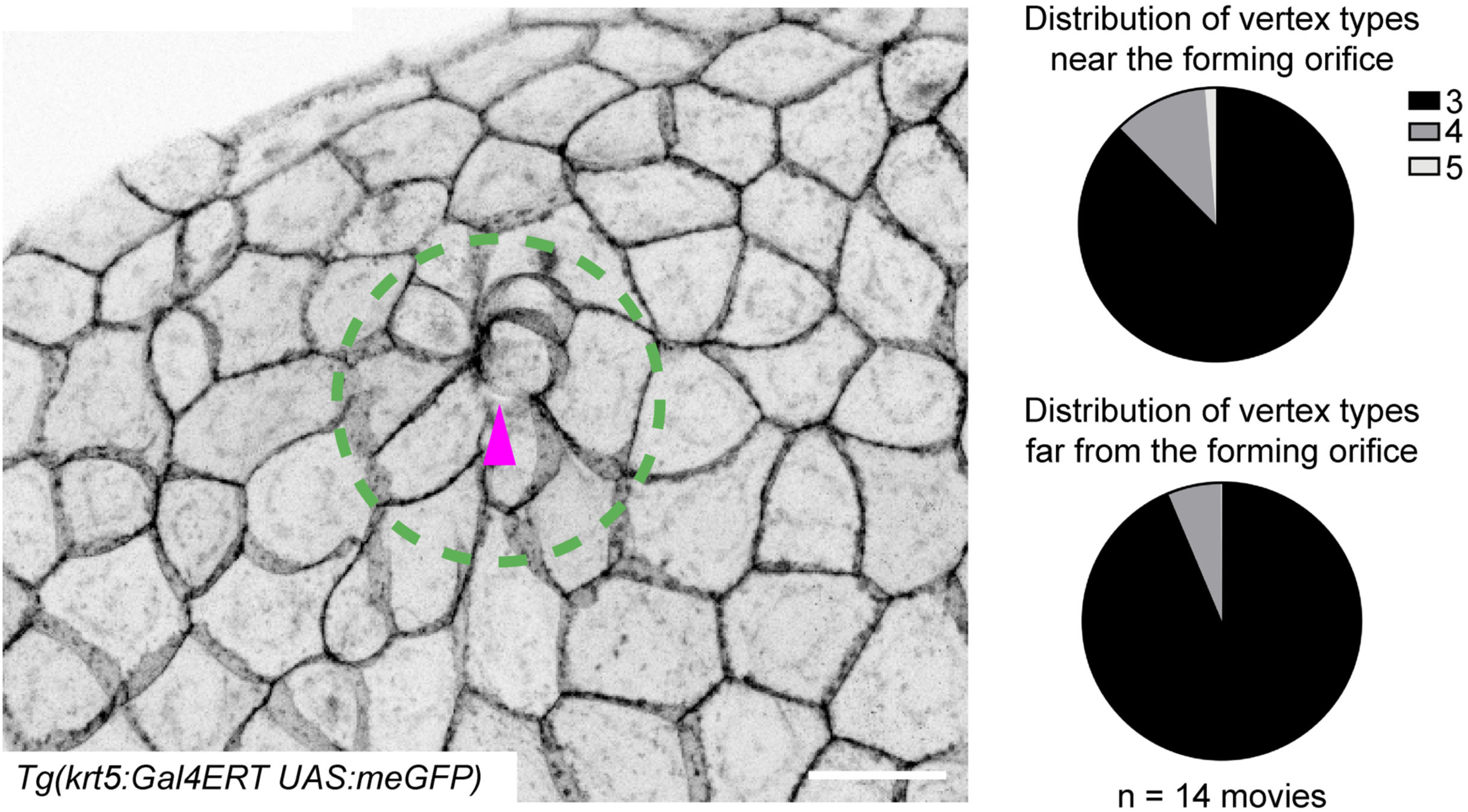
Analysis of vertex types in the skin epithelium. The different types of vertices (3, 4, 5 or more cells) were analysed on live embryos, in movies where the peridermal skin cells were labelled with membrane GFP, right at the moment of the orifice opening, as illustrated by the picture on the left. The magenta arrowhead points to the forming olfactory orifice. The two pie charts on the right present the distribution of vertex types (3, 4, 5 or more cells) in the skin peridermal layer covering the olfactory placode (at a distance < 25 µm from the opening orifice, within the green circle represented in the picture) and that of the skin located further away (at a distance > 25 µm from the opening orifice, outside of the green circle. Scale bar: 20 µm.

**Figure S2.**
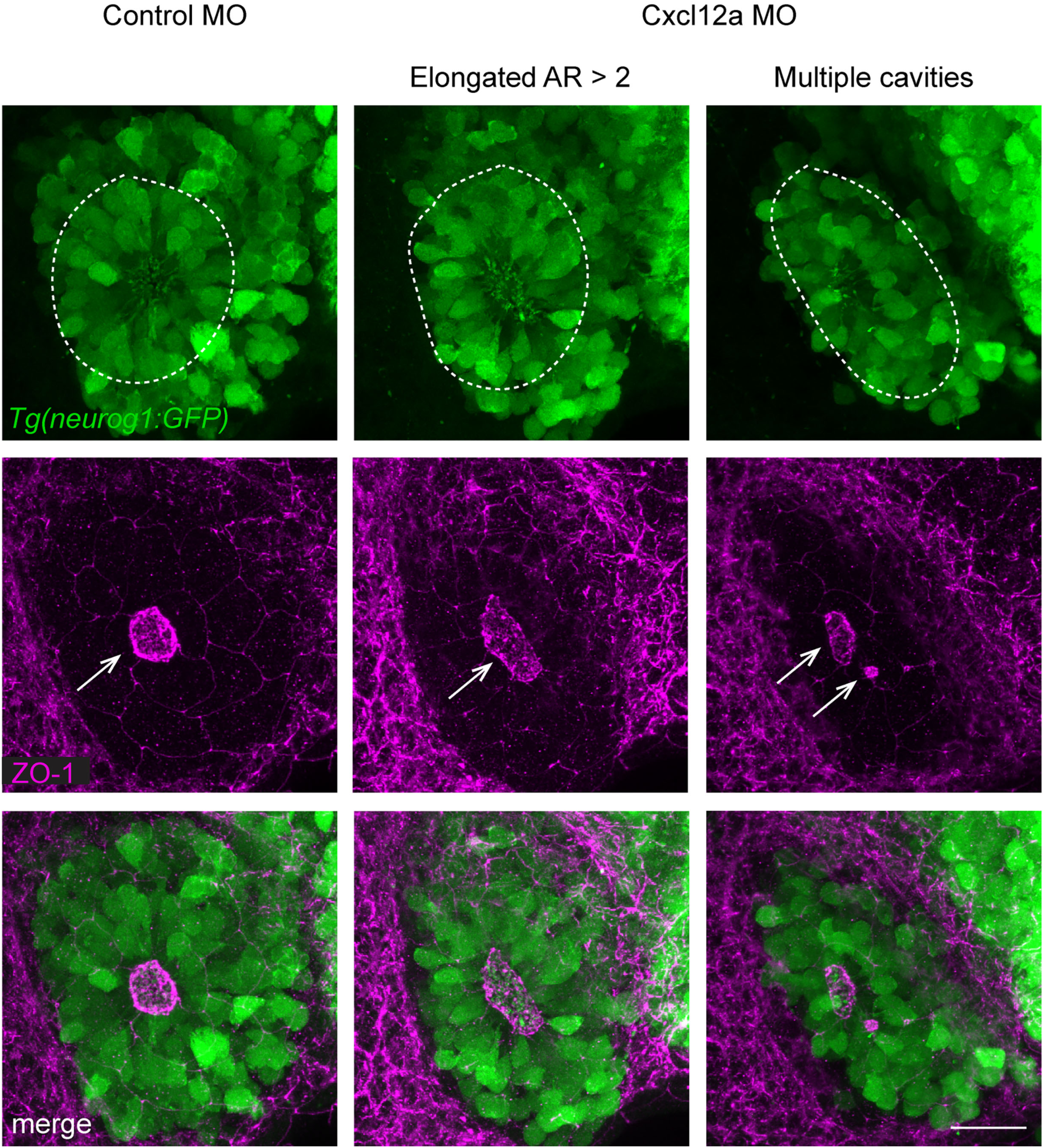
Organisation of the neuronal rosettes and of the rosette cavities upon Cxcl12a knockdown. Images illustrating the correspondence between the organisation of the neuronal rosettes, labelled with *Tg(neurog1:GFP)* (green), and that of the rosette cavities, labelled with ZO-1 immunostaining (magenta), in 34 hpf embryos injected with the control morpholino (left column) or the Cxcl12a morpholino (middle and right columns). The rosettes are surrounded by dotted lines. The arrows point to the rosette cavities. Scale bars: 20 µm.

**Figure S3.**
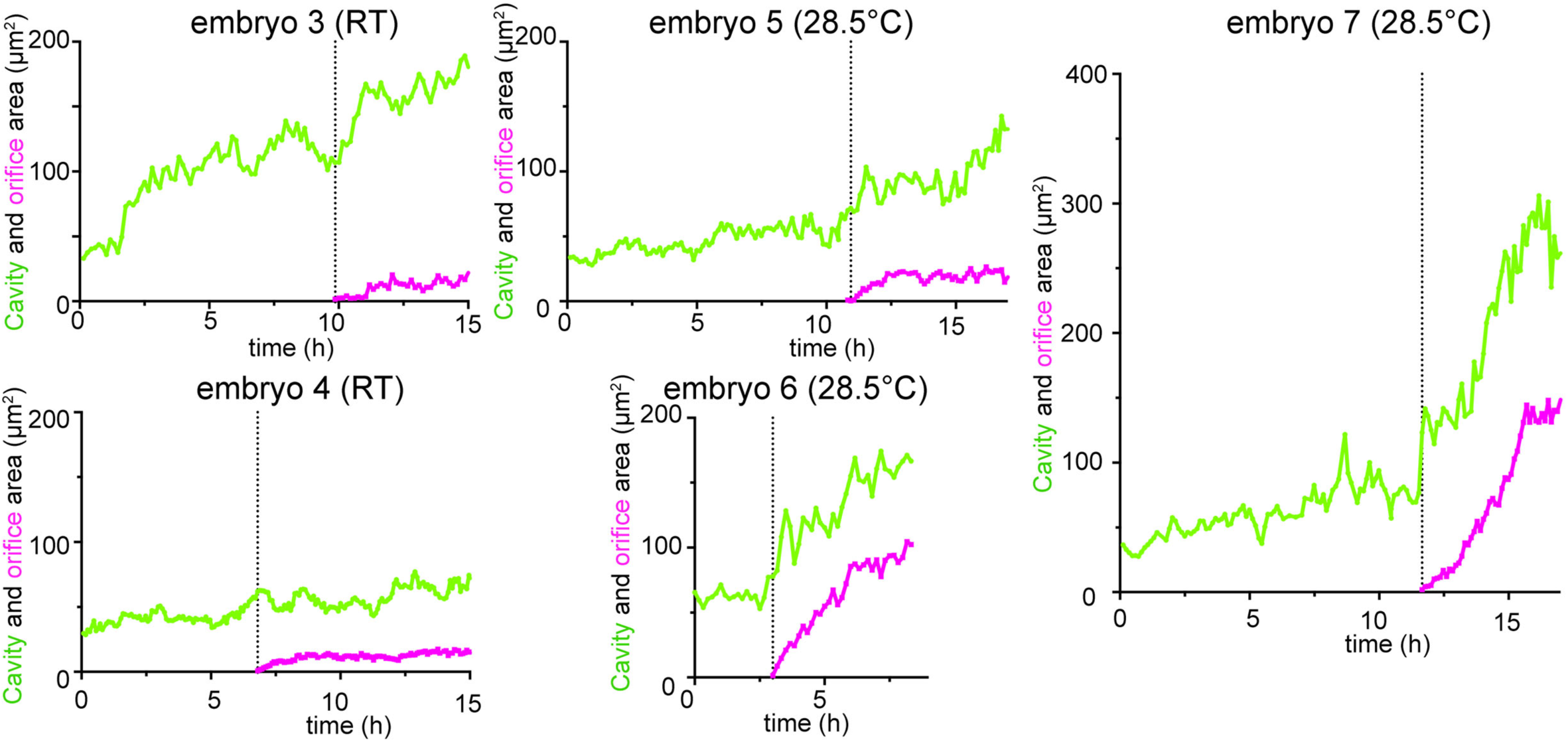
Quantification of the rosette cavity and orifice area as a function of time. Confocal live imaging was performed on *Tg(actb2:myl12.1-cherry); Tg(krt5:Gal4-ERT, UAS:meGFP*) embryos, *Tg(actb2:myl12.1-cherry); Tg(krt4:lynGFP)* embryos, or *Tg(actb2:myl12.1-eGFP); Tg(krt5:Gal4-ERT, UAS:RFP*) embryos to visualise myosin II dynamics and skin cell behaviours. The actomyosin cup was used as a readout for the rosette cavity, and its area was measured as a function of time. 4 independent experiments, of note two experiments were performed at room temperature (24°C) and the two others at 28.5°C. This figure presents the quantification for 4 embryos, the graphs for two additional embryos are shown in Figure 5D’.

**Figure S4.**
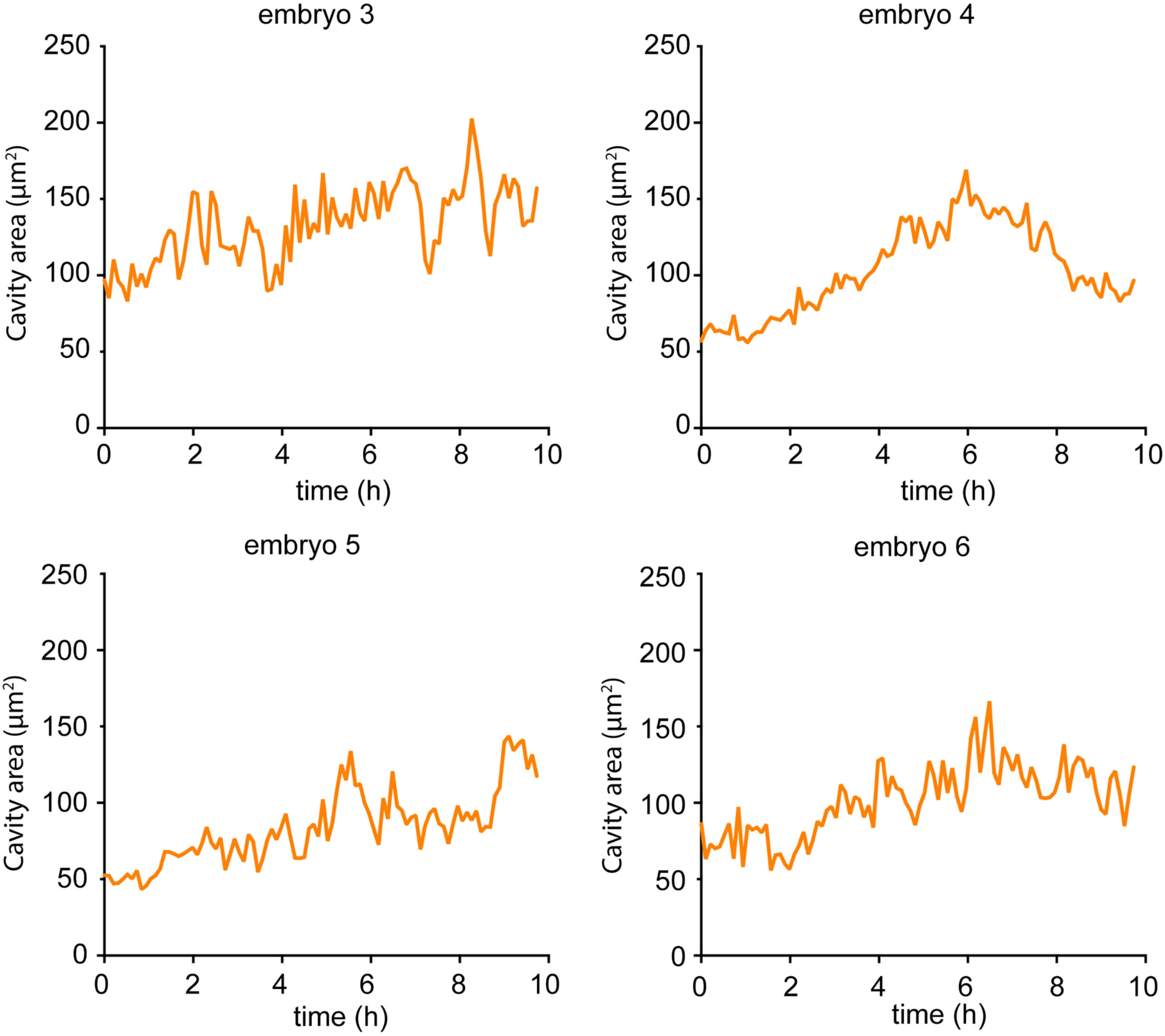
Quantification of the rosette cavity area as a function of time in blebbistatin-treated embryos. Confocal live imaging was performed on *Tg(krt5:Gal4-ERT, UAS:RFP*)*; Tg(actb2:myl12.1-eGFP)* embryos treated with blebbistatin from 26 hpf, in order to visualise myosin II dynamics and skin cell behaviours. The actomyosin cup was used as a readout for the rosette cavity, and its area was measured as a function of time. Note that in these embryos, the olfactory orifice does not open. This figure presents the quantification for 4 embryos, the graph for two additional embryos are shown in Figure 6B.

**Video S1. 3D stack showing the interaction between the olfactory neurons and the skin before the opening of the orifice** (same embryo as the z-section shown in Figure 1H). Confocal z-stack (dorsal view, the first z-sections are the most dorsal and the z-stack progressively goes ventral, anterior to the top, lateral on the left, z-step of 1 µm) on a fixed embryo at 29 hpf, just before the orifice opening, in which peridermal skin cells are labelled with *Tg(krt5:Gal4-ERT, UAS:RFP)* in magenta and all the early-born olfactory neurons are labelled with *Tg(neurog1:GFP)* in green. Note that upon fixation and for unknown reasons, the RFP signal is stronger in the nucleus that in the cytoplasm in some of the peridermal cells. The magenta asterisk indicates a peridermal cell which expresses the *UAS:RFP* transgene at very low levels, likely due to variegation. The most dorsolateral neurons of the placode are assembled into a rosette structure, and the apical tip of their dendrites is in close proximity with the overlying peridermal skin cells. The green asterisk indicates the hemispheric cavity present in the centre of the rosette and referred to as the rosette cavity. The white asterisks point to examples of peridermal cells that are deformed (rounded) in the vicinity of the future orifice, above the rosette cavity. The blue asterisk shows a peridermal cell sending a cytoplasmic protrusion towards the rosette cavity. Scale bar: 20 µm.

**Video S2. 3D stacks showing the interaction between the olfactory neurons and the skin before, during and after the opening of the orifice** (same embryos as the images shown in Figure 1I and Figure 3D). Confocal z-stacks (dorsal view, the first z-sections are the most dorsal and the z-stacks progressively go ventral, anterior to the top, lateral on the left, z-step of 1 µm) of *Tg(cldnb:meGFP)* embryos (cyan) fixed at 26, 28 and 34 hpf and immunostained for ZO-1 (yellow). The *Tg(cldnb:meGFP)* transgene labels the peridermal skin cells and all the cells of the olfactory placode, including the dorsal rosette and more ventral cells. The arrows point to the rosette cavities detected with ZO-1 accumulation at the apical tip of the rosette dendrites, and the arrowheads indicate the olfactory orifices at 28 and 34 hpf (at 26 hpf the skin has not opened yet). Scale bar: 20 µm.

**Video S3. Live imaging of skin cell behaviours during the opening of the olfactory orifice.** Confocal live imaging performed above the olfactory placode on a *Tg(krt5:Gal4-ERT, UAS:GFP)* embryo, in which peridermal skin cells are labelled with cytoplasmic GFP, showing that the olfactory orifice opens through the local loss of cell/cell contacts in the skin. See also Figure 2A. Maximum projection, dorsal view, anterior to the top and lateral to the left, delta t = 10 min. Scale bar: 20 µm.

**Video S4. Live imaging of skin cell behaviours during the opening of the olfactory orifice.** Confocal live imaging performed above the olfactory placode on a *Tg(krt5:Gal4-ERT, UAS:meGFP)* embryo, in which peridermal skin cells are labelled with membrane-bound GFP, showing that the olfactory orifice opens through the local loss of cell/cell contacts in the skin. The arrowheads point to the orifice when it starts opening. See also Figure 2B. Maximum projection, dorsal view, anterior to the top and lateral to the left, delta t = 12 min. Scale bar: 20 µm.

**Video S5. Live imaging of skin cells and placodal neurons during the opening of the olfactory orifice.** Confocal live imaging performed above the olfactory placode on a *Tg(krt5:Gal4-ERT, UAS:RFP); Tg(huC:eGFP)* embryo, in which peridermal skin cells are labelled in with RFP (magenta) and olfactory placode neurons are labelled with eGFP (green). Note that the orifice in the skin forms right above the centre of the placodal rosette. See also Figure 3A. Maximum projection, dorsal view, anterior to the top and lateral to the left, delta t = 12 min. The right panel shows the merge movie. Scale bar: 20 µm.

**Video S6. 3D stack showing a Cxcl12a morphant embryo in which two distinct rosette cavities are associated with two different holes in the skin** (same embryo as the images shown in Figure 4A5-C5). The embryo is at 34 hpf, carries the *Tg(cldnb:meGFP)* transgene to visualise peridermal skin cells and all the cells of the olfactory placode (in magenta), and has been immunostained for the tight junction protein ZO-1 (in green) to visualise the rosette cavities. Dorsal view, anterior to the top right and lateral to the top left, the first z-sections are the most dorsal and the z-stacks progressively go ventral, z-step of 1 µm. The right panel shows the merge image. The arrows point to the two rosette cavities and the arrowheads indicate the two olfactory orifices. Scale bar: 20 µm.

**Video S7. 3D stack showing actin distribution in the skin and in the placodal neurons before the opening onset** (same embryo as the images shown in Figure 5C, C’). The embryo is at 28 hpf, carries the *Tg(neurog1:KalTA4, UAS:meGFP)* transgenes to visualise the membrane of olfactory placode neurons (green) and has been stained with Rhodamine-phalloidin to visualise F-actin (magenta). The left panel shows the merge image, and the right panel shows the phalloidin staining alone. Dorsal view, anterior to the top, lateral on the left, the first z-sections are the most ventral and the z-stack progressively goes dorsal, z-step of 0.3 µm. The arrow indicates the accumulation of actin at the apical tip of the rosette dendrites, and arrowheads show examples of radial actin cables observed in placodal neurons. Scale bar: 20 µm.

**Video S8. High spatiotemporal resolution imaging of myosin II dynamics in the placode neurons right before the opening of the orifice.** Airyscan live imaging was performed on a *Tg(krt5:Gal4-ERT, UAS:RFP); Tg(actb2:myl12.1-eGFP)* embryo at 28 hpf injected with Cas9 and a GFP-targeted Crispr guide RNA to generate mosaic expression of myl12.1-eGFP. Maximum projection, dorsal view, anterior to the top and lateral to the left, delta t = 3 min 40 s. The left panel shows the merge movie, and the right panel shows the GFP labelling alone. The movie is shown twice. In the first round, arrowheads point to the static accumulation of myosin II at the apical extremities of the rosette dendrites. In the second round, examples of dynamic, radial myosin II cables are indicated by blue arrowheads in the first time point. Scale bar: 20 µm.

**Video S9. Live imaging of myosin II and skin cells during the opening of the olfactory orifice.** Confocal live imaging performed above the olfactory placode on a *Tg(krt4:lynGFP); Tg(actb2:myl12.1-mCherry)* embryo, in which peridermal skin cells are labelled in with membrane GFP (magenta) and myosin II dynamics can be visualised with ubiquitous *myl12.1-eGFP* expression (green). Arrowheads point to the forming orifice. Note that the actomyosin cup structure lining the rosette cavity expands exactly when the orifice opens in the skin. Arrows show examples of radial myosin II cables that appear to pull on and deform the actomyosin cup structure lining the rosette cavity, thus contributing to its expansion. Maximum projection, dorsal view, anterior to the top and lateral to the left, delta t = 530s. The right panel shows the merge movie. Scale bar: 20 µm.

